# Hyaluronan and CD44 targeting reverses early matrix changes, proinflammatory signals and fibrosis in primary sclerosing cholangitis

**DOI:** 10.64898/2026.07.21.739845

**Authors:** Vidushi Bansal, Lorand Vancza, Weiguo Fan, Koshi Kunimoto, Tzu Han Lo, Nhu Nguyen, Andrew Richardson, Antonios Chronopoulos, Rui Chen, Xingcai Zhang, Alexandre de Fraipont, Yuan Li, Yi Wei, Gregory W. Charville, Shuang Li, Nadine Nagy, Paul L Bollyky, Natalie J. Torok

## Abstract

Primary sclerosing cholangitis (PSC) is a rare, progressive liver disease characterized by biliary inflammation and bile duct strictures and no approved medical therapy. Despite its clinical severity, the pathological mechanisms underlying PSC remain poorly understood, largely due to early diagnostic challenges. Here we provide complementary evidence in human PSC samples, transcriptomic data, mouse models, and 3D cholangiocyte cultures that underscore the importance of hyaluronic acid (HA) and its cognate receptor CD44 in PSC pathogenesis. HA is a glycosaminoglycan abundant in the extracellular matrix in inflammatory disorders, yet its role in PSC has not been well characterized. We demonstrate that in early-stage PSC, cholangiocytes aberrantly produce high molecular weight HA that accumulates in the peribiliary matrix, increasing local tissue stiffness. This mechanical signal is transduced by a CD44/Integrin β1 receptor complex in cholangiocytes, driving cell proliferation, YAP mechanosignaling, pro-inflammatory cytokine production with a transition to a ductular reactive phenotype. CD44 knockdown in cholangiocyte cell lines and mouse models significantly lowered stiffness, and attenuated inflammation. Together, these findings reveal a mechano-inflammatory axis in which HA-driven matrix stiffening perpetuates biliary inflammation and disease progression, identifying HA targeting and CD44 as promising therapeutic strategies.

**One Sentence Summary:** Hyaluronan and CD44 mediate matrix changes and progressive fibrosis in primary sclerosing cholangitis.

## Introduction

Primary sclerosing cholangitis (PSC) is an idiopathic, cholestatic liver disease that leads to progressive inflammation and segmental fibrosis, complicated by recurrent cholangitis, and at later stages portal hypertension, cirrhosis, and cholangiocarcinoma. Although it is associated with inflammatory bowel disease, treating ulcerative colitis (UC) or Crohn’s has no major effect on PSC progression. Given the lack of effective pharmacotherapy, PSC is a major unmet medical need, with liver transplantation the only therapeutic option for select patients (*1*). PSC is usually diagnosed at advanced stages when clinical symptoms due to fibrosis, and stricturing arise, and as no potent anti-fibrotic therapy is available these patients are only treated symptomatically. The etiology and early pathomechanism of PSC still enigmatic given the lack of biological samples and models to study disease onset and early stages. In addition, changes in the peribiliary extracellular matrix, and the corresponding cholangiocyte behavior have not been well studied.

Hyaluronic acid (HA) is a major glycosaminoglycan in the extracellular matrix (ECM), consisting of D-glucuronic acid and N-acetyl-d-glucosamine repeats. It is synthesized as high molecular weight hyaluronan (HMW-HA) by the three hyaluronan synthases HAS1-3. It has a role in maintaining ECM architecture during development, and in certain instances it contributes to wound healing, fibrosis and inflammation (*2, 3*). According to current paradigms, low molecular weight hyaluronan (LMW-HA) is generated from HMW-HA following exposure to hyaluronidases, matrix metalloproteinases or reactive oxidant species (ROS) (*4*), (*5*), and has a proinflammatory role. LMW-HA can act as endogenous extracellular danger signal, promoting an inflammatory response trough TLR-2 (*6*), (*7*). HMW-HA in contrast, is thought to play a homeostatic/anti-inflammatory role (*8*) (*9*).

Along with its roles in inflammation, HA influences local mechanobiology. Chemically, the repeating disaccharide subunits of HMW-HA carry dense negative charges that attract counter-ions, generating a Donnan effect and higher osmotic swelling pressure. Interestingly, recent studies have shown that HMW-HA can stiffen the collagen network upon swelling, straightening and stretching the initially relaxed collagen fibers (*10*). Another study implicated HMW-HA in augmenting the compressive stiffness of self-assembled neocartilage by increasing the osmotic pressure and subsequently the resistance to compressive load (*11*).

As early matrix changes, mechanosensing, and the role of HA in disease progression have not been addressed, we focused on the peribiliary ECM, and CD44 as a cholangiocyte mechanoreceptor with a goal of developing new therapeutic approaches for PSC.

In this study we examine how early cholangiocyte HA production, preceding major collagen deposition, leads to increasing stiffness of the peribiliary matrix, and mechanocellular responses. We show that the CD44/integrin β1 complex has mechano-sensitive function, promoting cellular changes to an inflammatory reactive ductular phenotype. Deletion of cholangiocyte CD44 can abrogate early matrix stiffening due to HA in vivo. Inhibition of HA synthesis by 4-methylumbelliferone glucuronide (4-MUG) prevents stiffening of the matrix both at early and later phases of PSC, mitigating the expansion of ductular reactive cells (DRC) and subsequent inflammatory cascades. Together, these findings reveal a mechano-inflammatory axis in which HA-driven matrix stiffening perpetuates biliary inflammation and disease progression, identifying HA and CD44 as promising therapeutic targets in PSC.

## Results

### Cholangiocytes express CD44 and HA accumulates in the peribiliary matrix at an early stage of the disease

To evaluate the early changes in the peribiliary ECM prior to the onset of collagen deposition/fibrosis, we performed a time course study in the 3,5-diethoxycarbonyl-1,4-dihydrocollidine (DDC) dietary model that is known to develop cholangiopathy, ductular reaction and periductal fibrosis at later stages (*12*). We sacrificed mice after 48h or at 96h on this diet (Fig. 1A). The Mdr2^-/-^ model already exhibits peribiliary fibrosis (and therefore increased stiffness) at the time of birth. Therefore we elected to use the DDC model where early changes can be better discerned. (Fig. 1A). We observed that HA, visualized by hyaluronan binding protein (HABP) immunofluorescence (IF) started accumulating around CK19 positive bile duct cells only after 48h on the diet (Fig. 1B, C). Interestingly, hepatic stellate cell activation, visualized by collagen deposition (middle panel) and smooth muscle alpha actin (aSMA), lower panel), were not seen at this time point, but only later, after 96h on the diet (Fig. 1B), and Suppl 1A, individual channels, and Suppl 1B, antibody control).

**Figure 1.**
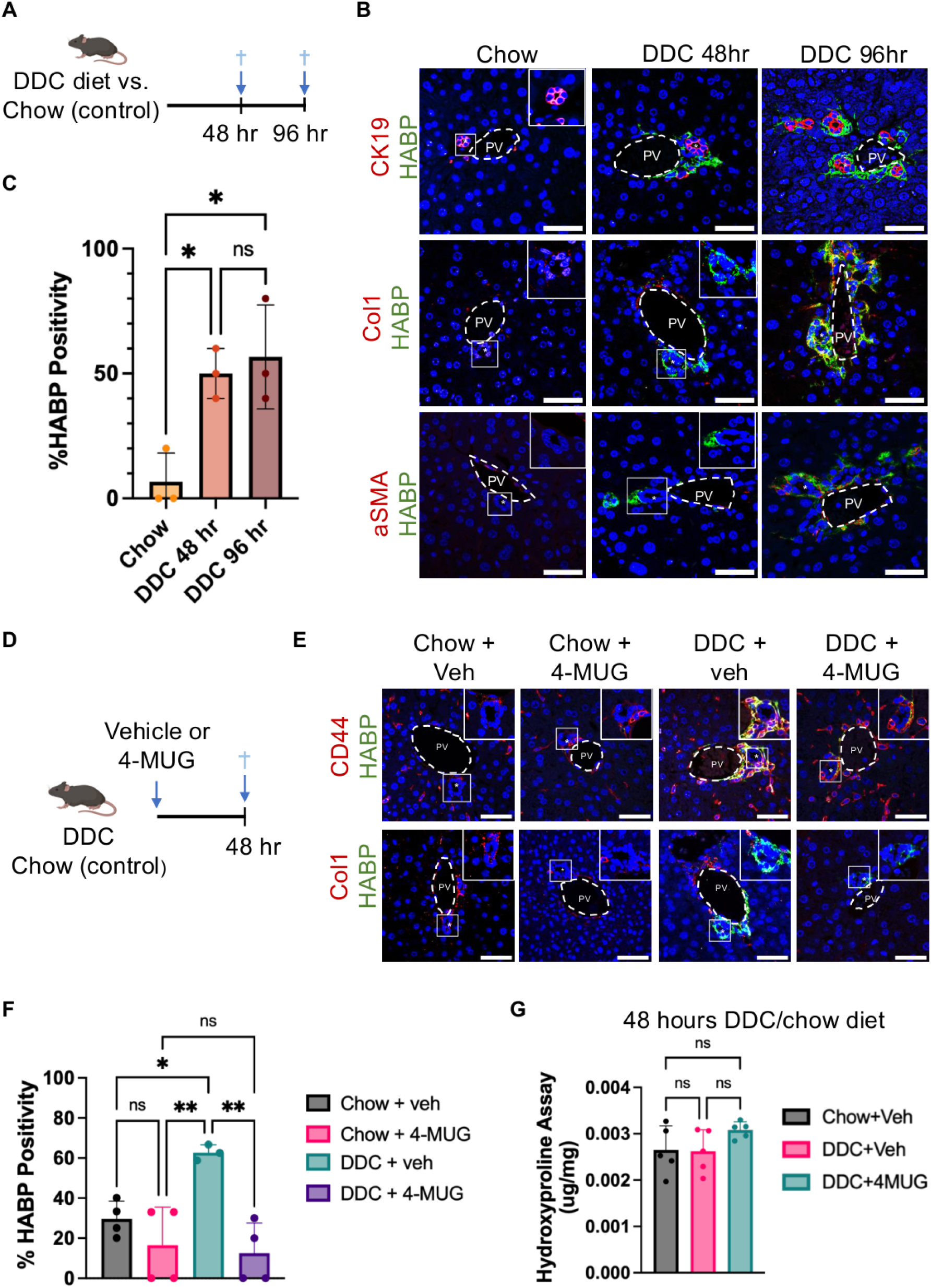
Early deposition of hyaluronan in the periductal areas in PSC. **(A)** Schematic of the early PSC model using the 3,5-diethoxycarbonyl-1,4-dihydrocollidine (DDC) vs. chow diets, and sacrificing mice after 48 or 96 hours on the diet (6-8 mice/group). **(B)** Representative immunofluorescence (IF) images using confocal microscopy depict representative merged images of hyaluronan binding protein, (HABP, green), and CK19 (magenta), HABP and collagen I (red), HABP and smooth muscle alpha actin (aSMA), as well as DAPI. (the separate HABP and collagen I images are shown in Suppl. Fig. 1a). Boxed areas and asterisks depict enlarged images of the bile ducts/periductal areas. **(C)** Percentage of hyaluronan binding protein (HABP) positive bile ducts in all conditions from B (N= 3 mice/group, 5 ROIs/each, mean±SEM, *P < 0.05, one-way ANOVA). **(D)** Schematic of the DDC mouse model using the HA synthase inhibitor 4-Methylumbelliferyl-b-D-glucuronide (4-MUG), vs vehicle. **(E)** Representative confocal IF images of HABP (green), CD44 (red), and collagen 1 (red) in mice on DDC/chow diet for 48 hours with 4-MUG or vehicle administration. **(F)** Percentage of hyaluronan binding protein (HABP) positive bile ducts in all conditions from E (N= 3 mice/group, 5 ROIs/each, mean±SEM, *P < 0.05, **P < 0.01, one-way ANOVA). **(G)** Hydroxyproline assay quantifying collagen in mice on chow and DDC diet, treated with 4-MUG vs. vehicle. Data are for n=5 animals/group, mean±SEM is shown, ordinary one-way ANOVA.

Next, we studied the effects of inhibiting hyaluronan synthesis using 4-MUG in this model (Fig. 1D). We decided to use 4-MUG rather than 4-Methylumbelliferone (4-MU) because the former molecule is more water soluble (greatly facilitating in vivo studies) and can suppress HAS without depleting the HA precursor UDP-glucuronic acid (*13*). We observed that HA deposition and cholangiocyte CD44 signal decreased following 4-MUG administration, in mice after 48h of DDC (Fig. 1E, F). At this early stage of the disease there was no significant induction in collagen production between the chow and DDC groups with either 4-MUG or vehicle treatment as assessed using a hydroxyproline assay (Fig. 1G), and picrosirius red staining (Suppl. Fig. 1C). Also, Sox9 positive ductular reactive cells decreased after 4-MUG treatment (Suppl. Fig. 2A, B). As early HA accumulation may affect the expansion of ductular reactive cells, we sought to examine the physical impact of these changes on the extracellular matrix.

### HA deposition elicits stiffening in the peribiliary ECM

We next analyzed changes in the mechanical properties of the matrix, using atomic force microscopy (AFM) on ex vivo liver sections (Fig. 2A). We tested the peribiliary and more distant central parenchymal areas and used these data to generate a heatmap of Young’s moduli, denoting stiffness. We observed that peribiliary stiffness was already increased after 48h and this was reversed in the DDC group by HA synthesis inhibition using 4-MUG treatment (Fig. 2B, C). Testing the central areas (zone 3) on corresponding samples, there was no significant difference in stiffness between the groups (Fig. 2D). However, there was overall a lower dissipated energy (viscoelasticity) in the periportal areas on DDC diet (Suppl. Fig. 3A) whereas no significant change was seen in the central areas (Suppl. Fig. 3B).

**Figure 2.**
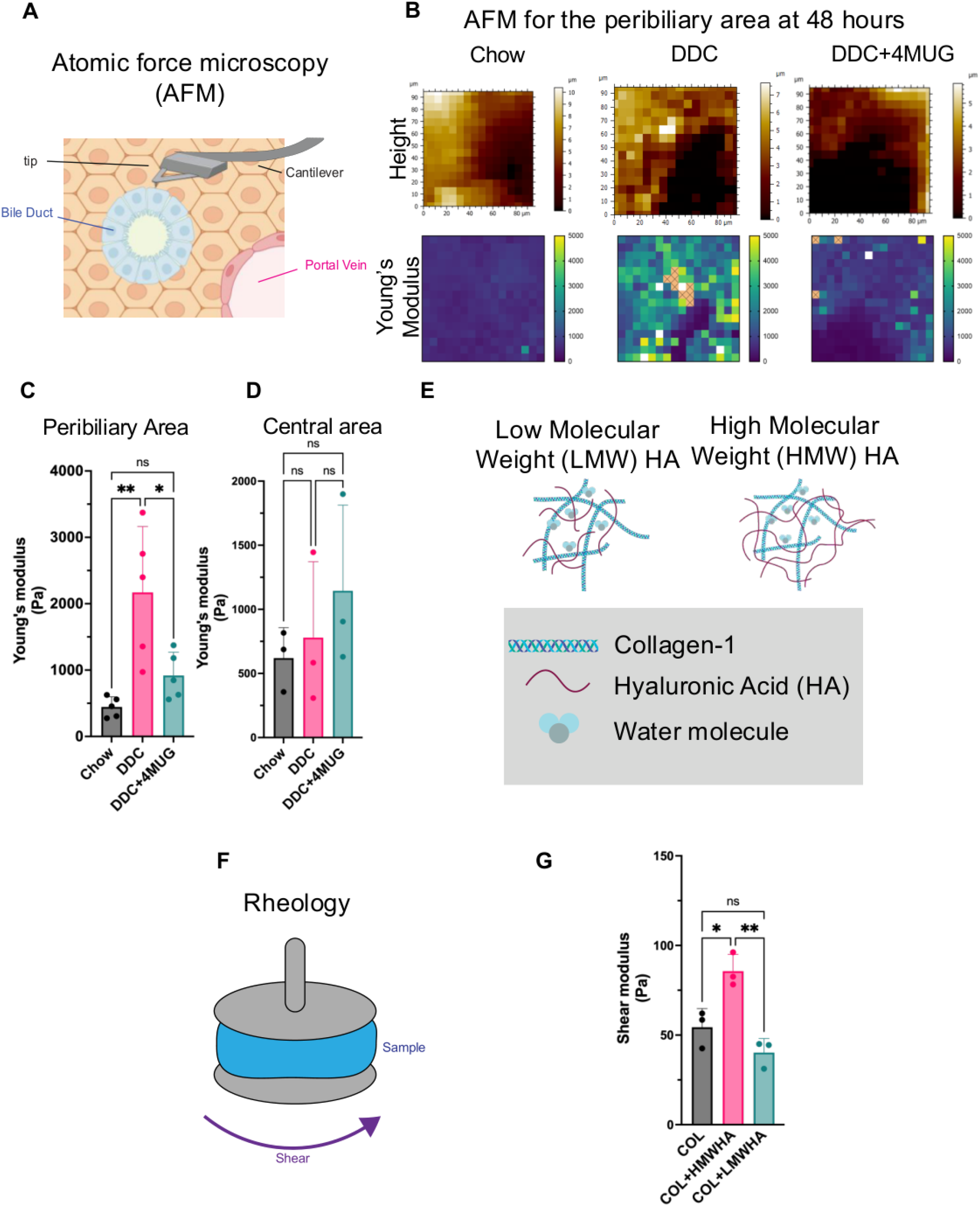
Accumulation of hyaluronic acid increases matrix stiffness in the peribiliary area. **(A)** Schematic of the atomic force microscopy, testing the peribiliary or pericentral areas, using indentation. Multiple (5 measurements) areas were indented. **(B)** Representative DAPI images boxed area for the peribiliary areas, show the tip of the probe, the height sensor images and the Young’s modulus heatmaps for mice on chow and DDC diet for 48 hours with 4-MUG or vehicle treatment. **(C)** The Young’s moduli data are depicted for the peribiliary, and **(D)** central areas. *P < 0.05, **P < 0.01, n=5/group, 3 ROIs/mouse; and n=3/group 3 ROIs/mouse (respectively), mean±SEM, ordinary one-way ANOVA. **(E)** Schematic of the telopeptide intact collagen I-hyaluronic acid hydrogel. 3D hydrogels were constructed using collagen I mixed with either low or high molecular weight hyaluronic acid. **(F)** Schematic of shear rheology used to test the physical properties of the hydrogels. **(G)** Mean shear moduli are shown for pure collagen I, collagen-1 + high molecular weight HA (HMWA), and collagen I + low molecular weight HA (LMWHA) hydrogels. *P < 0.05, **P < 0.01, data are for n=3, and were analyzed using a one-way ANOVA, mean±SEM is shown.

To further study the mechanical role of HA, we generated 3D hydrogels with collagen I and with either HMW-HA or LMW-HA (Fig. 2E) and tested the mechanical properties by rheometry by strain sweeps with increasing strain amplitudes after 24h of swelling (Fig. 2F, G). The shear modulus significantly increased when collagen was mixed with HMW-HA compared to the pure collagen gel, but this was not seen with LMW-HA (Fig. 2G). Notably, the loss modulus did not change significantly (Suppl. Fig. 3C).

Together, these results suggest that the peribiliary matrix exhibits stiffening in the early stages of PSC due to HMW-HA accumulation.

### Stiff ECM promotes a reactive cholangiocyte phenotype through CD44 and integrin **β**1 mediated mechano-signaling

To directly test the mechanical effect of peribiliary stiffness on cholangiocytes, we embedded H69 cells in collagen-alginate interpenetrating network (IPN) hydrogels, with tunable stiffness (Fig. 3A). By adjusting the Ca^2+^ concentration, the hydrogel stiffness can be tuned. We used soft and stiffer conditions with a Young’s modulus of 1 kPa (representing the physiological softer matrix), and 2.5 kPa, respectively (*14*). The stiffer ECM promoted cholangiocyte proliferation (Ki67), and the expression of the HA receptor CD44, IL-6, and MCP-1 (Fig. 3B), as well as increase in HAS1 and 2 (Fig. 3E). Previously these were the two main HAS enzymes that showed an induction in the PSC model whereas HAS3 did not change (*15*). We also detected stronger HAS2 and HABP signals colocalizing in stiffer matrices (Fig. 3C), as HAS2 is known to extrude HA directly into the ECM (*16*).

**Figure 3.**
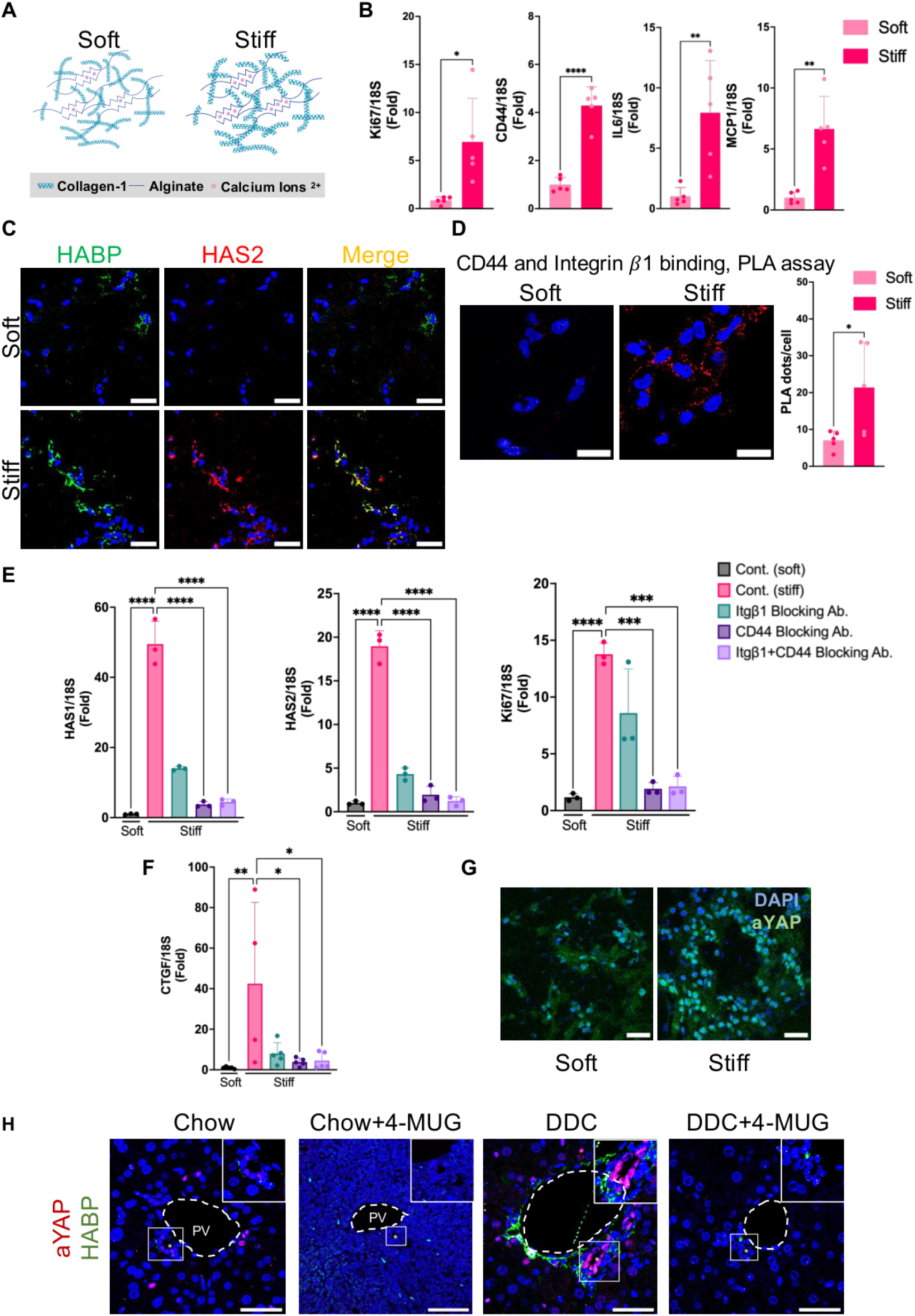
CD44 acts as a mechanosensor in cholangiocytes. **(A**) Schematic of soft and stiff 3D Collagen-1 and Alginate iPN hydrogels (CAiPNs). **(B)** RTqPCR of Ki67, CD44, IL-6, and MCP-1 mRNA expression in H69 cells in soft vs. stiff CAiPNs. (*P < 0.05, **P<0.01, ****P<0.001, n=5, mean±SEM, Unpaired t-test). **(C)** Representative images of H69 cells immunostained for HABP (green), HAS2 (red), nuclei (blue) in soft vs. stiff CAiPNs (bar= 20 um). **(D**) Proximity ligation assay (PLA) depicts direct binding between CD44 and integrin β1 (ITGB1) in soft vs stiff CAiPNs (scale bar 25 μm). The PLA signal was analyzed (30 cells in 5 gels per group, n = 5, each, *P < 0.05, Anova). **(E)** HAS1, HAS2, and Ki67 mRNA expression levels as assessed by RTqPCR in H69 cells after using blocking antibodies to CD44 and/or integrin β1 (ITGB1). ***P<0.005, ****P<0.001, n=3, mean±SEM, one-way ANOVA). **(F)** YAP target CTGF mRNA studied by RTqPCR in H69 cells after blocking CD44 and/or integrin β1 (ITGB1), in stiff CAiPNs. (**G)** Active nuclear YAP (green) is depicted in H69 cells in soft or stiff matrices, in representative immunofluorescence images. **(H)** Representative confocal IF images of active YAP in cholangiocyte nuclei and HABP in mice on DDC diet with 4-MUG vs. vehicle treatment for 48 hours (scale bar 20 um).

To evaluate the mechanosensory role of CD44, we hypothesized that it might structurally cooperate with integrin β1 (ITGB1), a known sensor that engages the mechanical clutch upon higher ECM stiffness. By proximity ligation assays, we found increased binding of CD44 to ITGB1 when matrix stiffness increased (Fig. 3D, Suppl. Fig. 4). Either antibody-mediated inhibition of CD44 or integrin β1, or both, significantly reduced HAS1 and HAS2 expression, as well as cholangiocyte proliferation (Fig. 3E). Stiffness also induced the expression of CTGF in a CD44 and/or ITGB1-dependent manner, pointing to the activation of the Hippo/YAP pathway (Fig. 3F). Indeed, we found active nuclear YAP in cholangiocytes cultured in stiff 3D matrices (Fig. 3G). We similarly observed active YAP in vivo in the DDC model after 48 hours and found that this this was inhibited by the HAS inhibitor, 4-MUG (Fig. 3H).

To study the CD44 receptor domains involved in mechano-transduction, we transfected H69 cells either with the wild type plasmid, or a version of CD44 mutated at R41A, which disrupts interactions between Arginine R41 and the polysaccharide chain of HA, thus inhibiting CD44-HA interactions (*17*), (Fig. 4A). We observed that there was a significant reduction in proliferation of cholangiocytes in stiff matrices, HAS2 and CTGF mRNA expression, and the production of proinflammatory markers IL-1β and TNF-α (Fig. 4B). These data point to a CD44-driven molecular circuit whereby stiffening of the peribiliary matrix drives cholangiocyte proliferation and transition to a ductular reactive phenotype via CD44/integrin β1 mechano- signals.

**Figure 4.**
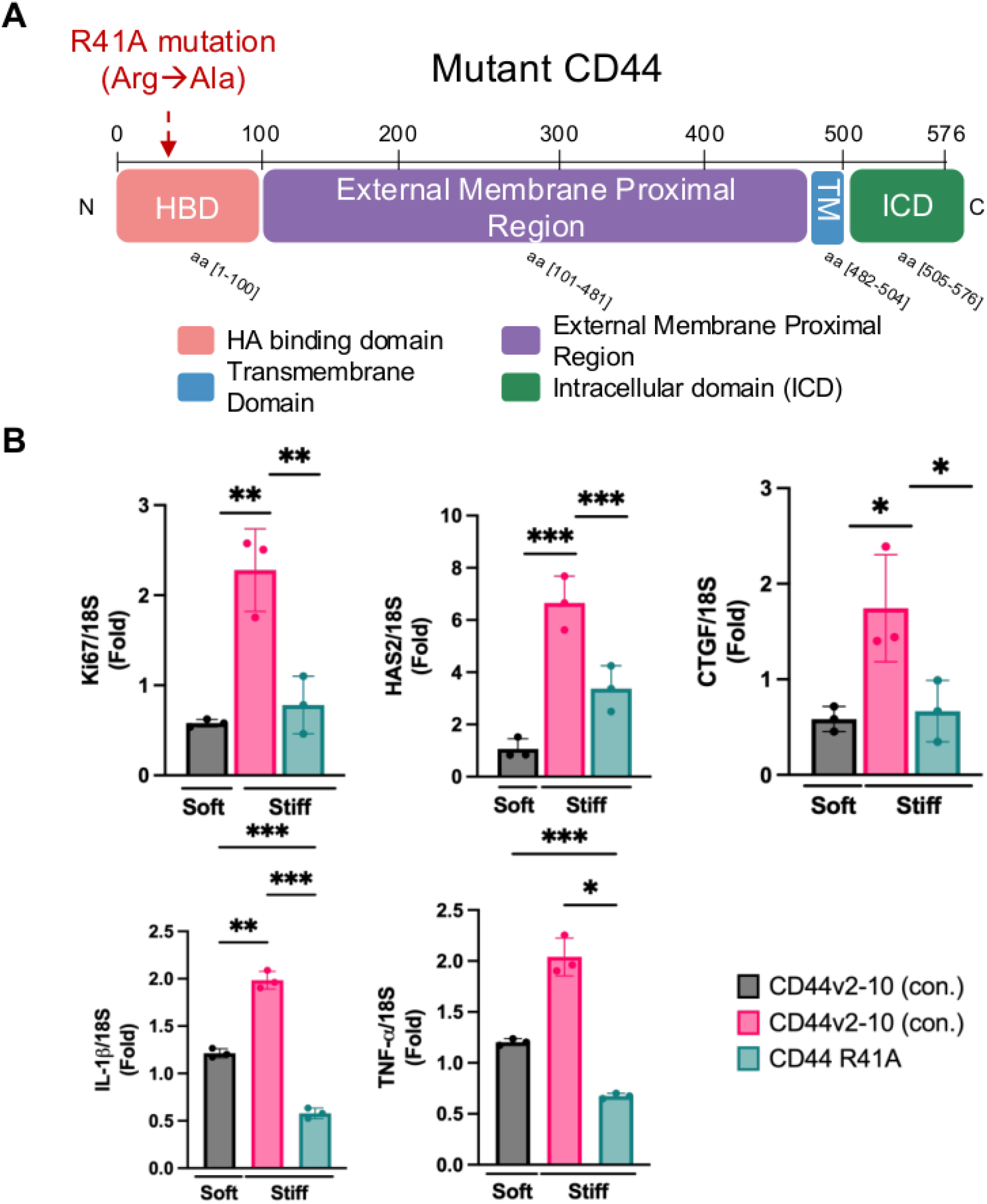
CD44 receptor mutant reduces cholangiocyte proliferation, the expression of HAS2, and proinflammatory cytokines in stiff matrices. (**A**) Schematic of human CD44v2-10 receptor domain organization and point mutation (CD44^R41A^) in the extracellular domain that prevents binding of HA. (**B**) The expression of Ki67, HAS2, CTGF, IL-1β, and TNF-α mRNA, assessed by RT-qPCR in cells transfected by CD44v2-10 (control) or CD44^R41A^ and cultured in stiff CAiPNs. *P < 0.05, **P< 0.01, ***P< 005; n=3, mean±SEM, Welch test.

### Cholangiocyte CD44 KO mice exhibit reduced HA production, peribiliary stiffness and inflammation

To better understand the role of the HA receptor CD44 in intrahepatic cholangiocytes in vivo, we crossed CD44^fl/fl^ C57BL/6J mice with tamoxifen inducible Opn^CreER^ mice (Fig. 5A). The resulting strain was then backcrossed to a C57BL/6J background. CD44^fl/fl^ mice were used as controls. Mice were placed on DDC vs. chow diet for 48 hours. We observed that tamoxifen administration induced CD44 deletion from cholangiocytes (Fig. 5B), reduced peribiliary HA deposition (HABP, Fig. 5B and C), and nuclear active YAP, respectively, at 48 hours. We did not observe a difference in the low picrosirius red staining signal (Suppl. Fig. 5A) or change in aSMA positive myofibroblasts/portal fibroblasts (Suppl. Fig. 5B) at this early time point, indicating that fibrosis was not yet affected.

**Figure 5.**
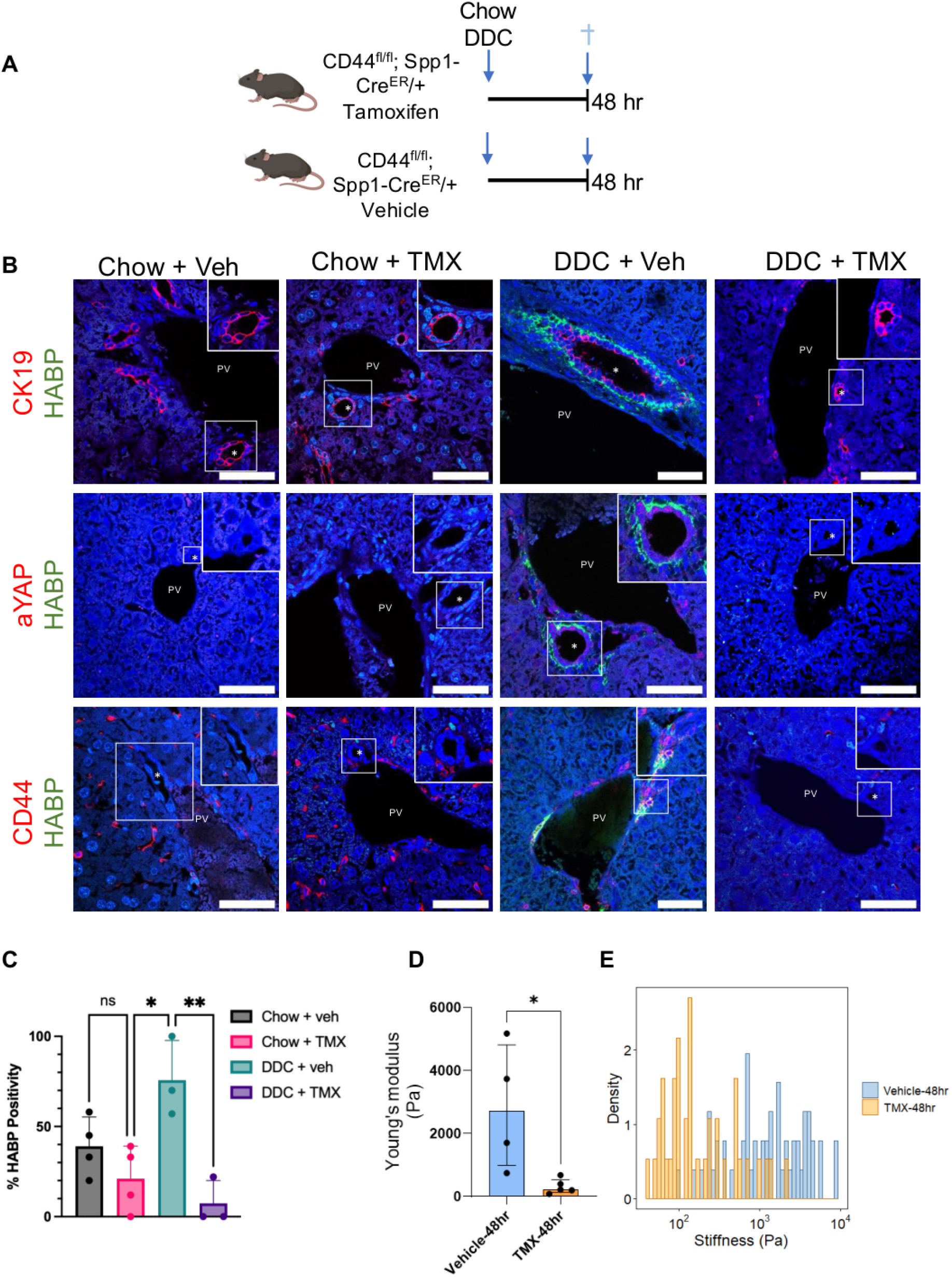
Cholangiocyte CD44 knockout model exhibits lower HA deposition and softening of the periductal matrix. **(A)** Schematic of the CD44 cholangiocyte KO (CD44^fl/fl^ x Opn^CreER/+)^ mouse model. Tamoxifen vs. vehicle were administered, then mice were on DDC or chow diets for 48hours (7-8 mice group). **(B)** Representative confocal IF images of HABP, combined with CK19, aYAP, or CD44. Asterisks depict the bile ducts, (boxed enlarged area), PV: portal vein, bar= 50 um. **(C)** HABP signal was assessed in 3-4 portal areas in 3 mice by ImageJ. *P < 0.05, **P <0.01; n=3, mean±SEM, one-way ANOVA. **(D)** Young’s moduli were measured by AFM in the peribiliary areas in mice on the DDC diet for 48h treated with vehicle vs. tamoxifen. *P < 0.05, n=4-5/group, 3 ROIs/mouse; mean±SEM, Welch’s t-test **(E)** depicts density plots of stiffness measurements.

Studying tissue sections from these mice and controls by AFM, we found that upon CD44 deletion from cholangiocytes the peribiliary area was much softer (Fig. 5D, E). In summary, these findings corroborate that in early PSC model, the periductal stiffening is driven by CD44/HA feedback loop.

### Fibrosis and inflammation improve after treating *Mdr2^-/-^*mice with the HA synthesis inhibitor 4-MUG

To evaluate the effects of the HA synthesis inhibitor 4-MUG at the later stages of the disease when fibrosis is already present, and a different model, we studied the *Mdr2^-/-^* mouse model (Fig. 6A). In this genetic PSC model fibrosis appears after birth, and transdifferentiated myofibroblasts are an important source of HA production in the liver (*15*). In these mice, we observed that increased HA accumulation was present around the bile ducts by week 8 (Fig. 6B). This was reduced after 4 weeks of 4-MUG treatment that begun at 4 weeks (Fig. 6A, B). Picrosirius red staining revealed reduced fibrotic areas in the *Mdr2^-/-^*mice treated with 4-MUG, compared to the vehicle-treated group (Fig. 6B, C), and reduced expression of procollagen α1(I), (Fig. 6D). Thinner collagen bands characterized by lower density were seen on second harmonic generation microscopy images (Fig. 6E). 4-MUG treatment significantly decreased CD44, as well as IL1-β, IL-6 and MCP-1 pro-inflammatory transcripts (Fig. 6F), and resulted in a lower number of active YAP positive cholangiocytes (Figs. 6G, H, and Suppl. Fig. 6, individual channels).

**Figure 6.**
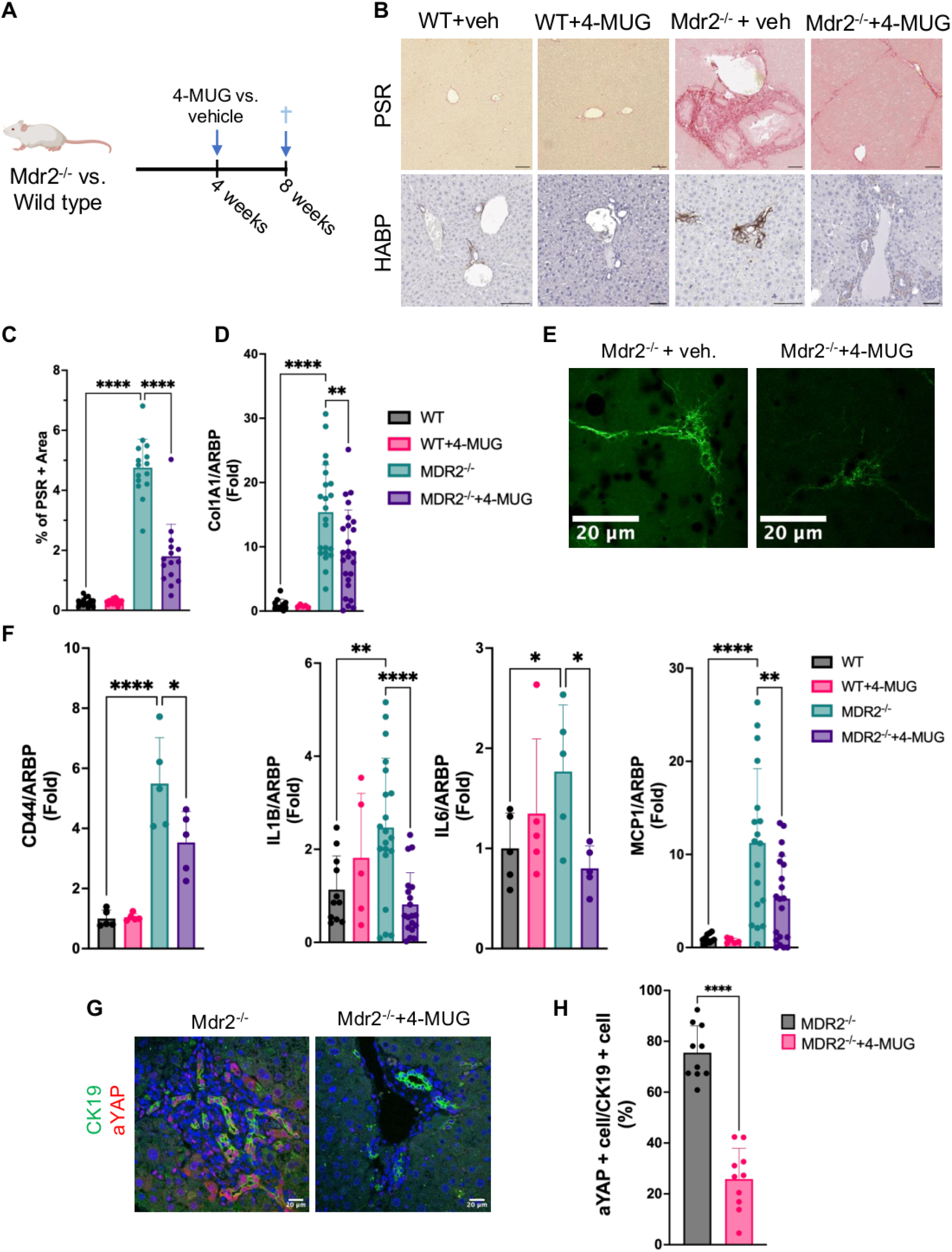
Hyaluronan synthase inhibitor 4-MUG improves fibrosis in a late PSC model. **(A)** Schematic of the Mdr2 ^-/-^ mouse model, with 4-MUG vs. vehicle treatment, therapeutically from week 4 to 8. **(B)** Representative picrosirius red images (top raw, bar=100um), and immunostaining for HABP (bottom panel, bar=100 um) in all experimental arms. **(C)** 4-MUG treatment significantly lowered picrosirius positive areas (****P < 0.001, 5 FOVs, n=3 , mean±SEM, one-way ANOVA), and **(D)** procollagen α 1(I) expression, **p< 0.01, ****P < 0.001, n=5-15 , mean±SEM, one-way ANOVA. **(E)** Representative second harmonic generation and two photon microscopy images depict collagen bands in vehicle vs 4-MUG treated Mdr2^-/-^ mice. **(F)** Expression of CD44, IL-1β, IL-6 and MCP-1 studied by RTqPCR in the different animal groups *P < 0.05, **P < 0.01, ****P < 0.001, n=5-15, mean±SEM, one-way ANOVA. **(G)** Representative immunofluorescence confocal microscopy of CK19 and active YAP in the vehicle vs. 4-MUG-treated mice, and **(H)** the percentage of aYAP + nuclei in CK19 + cholangiocytes in these mice, ****P < 0.001, n=4, mean±SEM, Welch’s t-test.

Taken together, these data demonstrate that in advanced PSC, 4-MUG mediated inhibition of HA accumulation results in an improvement in the expansion of ductular reactive cells, inflammatory responses, and fibrosis.

### CD44 and hyaluronan synthesis pathways are enriched in patients with PSC

We then performed immunohistochemistry on deidentified healthy or PSC patient liver samples obtained from the Stanford Biobank. We observed that CD44 and HABP signals were enriched in bile ducts/peribiliary areas (Fig. 7A, secondary only control, Suppl. 7).

**Figure 7.**
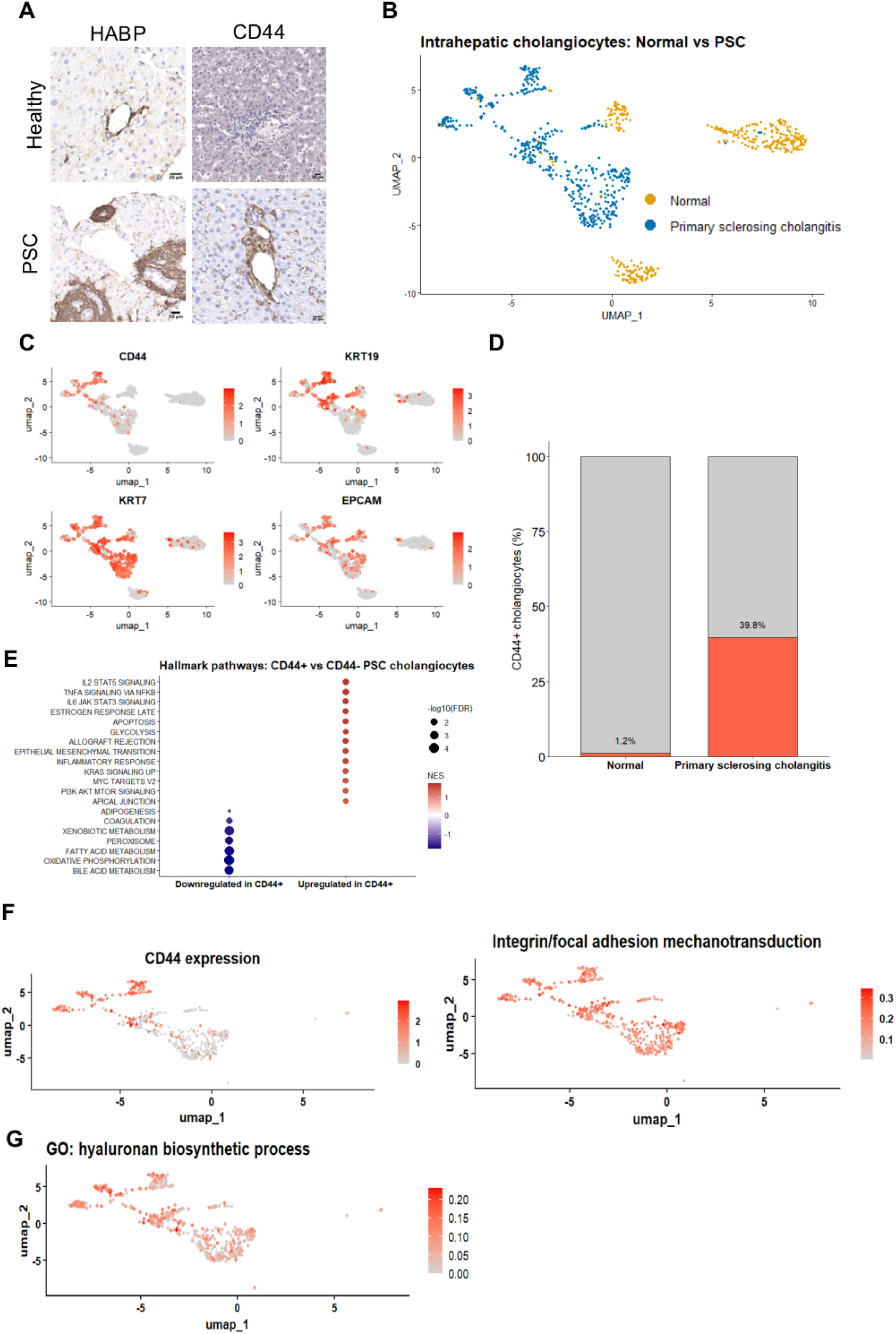
PSC patients express CD44 in cholangiocytes, and exhibit enrichment in HA synthesis pathways and integrin/focal adhesion mechanotransduction signatures. **(A)** Representative IHC images of CD44 and HABP in healthy and PSC patient livers. **(B)** UMAP projection of intrahepatic cholangiocytes from healthy and PSC patient livers (GSE247128) shows segregation of normal versus PSC cells. (**C)** FeaturePlots demonstrate preferential enrichment of CD44 in the PSC cholangiocyte cluster. (**D)** Quantification of CD44 transcript–positive cholangiocytes in PSC (n = 8 patients) compared with normal donors (n = 6). (**E)** Hallmark gene-set enrichment analysis comparing CD44+ versus CD44- PSC cholangiocytes. **(F)** Enriched CD44-high PSC patient cholangiocyte regions on the UMAP. These cells exhibit higher integrin/focal adhesion mechanotransduction signatures (AUCell scoring), and **(G)** enrichment of “hyaluronan biosynthetic processes”.

Next, UMAP projection of intrahepatic cholangiocytes from healthy and PSC livers (GSE247128) was done and this revealed segregation of normal versus PSC cells (Fig. 7B). FeaturePlots demonstrate preferential enrichment of CD44 in the CK19+ PSC cholangiocyte cluster (Fig. 7C), and a marked expansion of CD44^+^ cholangiocytes in PSC (n = 8 patients) compared with normal donors (n = 6 patients, Fig. 7D). Hallmark gene-set enrichment analysis shows induction of inflammatory programs in CD44-positive cholangiocytes compared to the CD44-negative cells (Fig. 7E). CD44 positive PSC cholangiocytes exhibit significantly higher integrin/focal adhesion mechanotransduction signatures based on AUCell-based module scoring (Fig. 7F), and elevated Gene Ontology scores for “hyaluronan biosynthetic process (GO:0030213)” compared to CD44 negative cells (Fig. 7G).

Together, these results indicate that CD44 and hyaluronan synthesis pathways are enriched in PSC patients.

## Discussion

Diagnosing and studying early stages of PSC in patients has been limited due to inaccurate diagnostic and imaging approaches, and scarcity of samples. In this study we aimed at modeling both earlier and later stage disease, and showed that cholangiocytes aberrantly produce HA before the onset of fibrosis and that it accumulates in the peribiliary matrix, increasing local tissue stiffness. This mechanical signal is transduced by a CD44/Integrin β1 receptor complex on cholangiocytes, promoting a cascade that can culminate in the recruitment of inflammatory cells, myofibroblast activation, and fibrosis. Conversely, inhibiting HA synthases potently reverses peribiliary stiffening and subsequent fibrosis. CD44 is a key to this pathological feed-forward signaling loop, as demonstrated by our findings that knockdown of CD44 in cholangiocyte cell lines and mouse models significantly attenuates inflammatory markers. CD44 is known to be highly expressed in the intrahepatic biliary epithelial cells in PSC patients (*18*) potentially linked to systemic autoimmune or inflammatory profiles (*19*). However, neither the mechanistic role of CD44 or its ligand HA have been addressed, and the early events in the peribiliary ECM have not been well understood.

Our findings reveal that HA deposition caused strain stiffening in the peribiliary ECM, and this was sensed by cholangiocytes via CD44 and integrin β1-mediated mechano-signaling promoting proliferation and a transition to a ductular reactive phenotype. These responses relied on intact extracellular signaling and HA binding domains and were augmented by complexing with integrin β1. Cooperation between integrin β1 and CD44 has been documented in other contexts such as in cancer cell adhesion (*20*), and CD44 clustering is associated with lipid raft reorganization and integrin β1-mediated cytoskeletal events (*21*). We saw that in stiffer matrices, there was CD44-integrin binding mediated Hippo/YAP activity which led to an increase in CTGF, and to a proinflammatory phenotype of cholangiocytes. These early changes elicited by mechanosignaling however, were independent of myofibroblast activation, or collagen deposition. This is clinically relevant as the cause(s) of segmental fibrosis in PSC are not well understood.

CD44 in different systems was shown to mediate stiffness-related mechano-signals (*10*),(*22*), (*23*),For instance, in autoimmune insulitis CD44 was associated with faster disease progression (*24*), CD44’s mechanosensitive function could potentially be explained by undergoing force-dependent conformational changes and switching between low affinity and high affinity HA-binding states. Its binding to integrin β1 may reinforce downstream actin mediated signals through the Ezrin, Moesin, or Radixin (ERM) family proteins. We observed a feedforward loop in vivo where deletion of CD44 led to lower HAS1/2 activity and HA production, resulting in matrix softening. This is consistent with previous work in other disease contexts such as glioblastoma and breast cancer where the presence of a positive feedback loop between CD44 and HAS2 was described (*25, 26*). In fact, these studies showed that upon CD44 knockdown, HA secretion also decreased through a downregulated HAS2 activity. The CD44–HAS–HA loop thus may contribute to the persistence and progression of biliary injury even after the initial trigger has diminished, though further studies will be needed to prove this clinically.

A new aspect of peribiliary HA deposition is that mainly HMW-HA but not LMW-HA deposition increased strain stiffening. Cells produce HMW-HA forming an intricate 3D structure that results in an increased viscosity of HA, reflective of greater entanglement (*27*). Conversely, LMW-HA—typically generated via enzymatic depolymerization by hyaluronidases—could potentially be present at later stages of PSC, and closely linked to ongoing inflammatory activity (*28*). While HMW HA is thought to have a protective role in cancer (*29, 30*), the mechanical consequences of its accumulation could be context and disease dependent. Elucidating the contributions of HA of different sizes in these responses will require further work, especially in regards to biliary diseases.

An intriguing question is how cholangiocytes regulate HA synthesis. Altered bile acid composition and ongoing exposure to gut derived microbial metabolites are conceivable signals. In inflammatory bowel disease, HAS2-mediated HA deposition was linked to non-responsiveness to infliximab and vedolizumab therapy in patients (*31*), and HAS inhibition conferred robust protection against murine DSS-induced colitis. We found that HA’s cognate receptor CD44 was also markedly induced already at the early stages of the disease. Further work is needed to define the key mediators, and studies on the induction of adaptive immune responses that may play a role in modulating the HAS/HA/CD44 axis. By disrupting the reciprocal interactions between ECM remodeling, cholangiocyte mechano-sensing, and immune activation, interventions targeting the HAS/HA/CD44 pathway may slow down disease progression.

We saw that in the fibrotic Mdr2^-/-^ mice, 4-MUG effectively decreased HA production, and lowered proliferation and cholangiocyte YAP activation culminating in protection against inflammation and fibrosis. We would like to emphasize that at this later stage, HA production is likely to be a cumulative effect of active myofibroblasts (*15*), and cholangiocytes that have transitioned to a ductular reactive phenotype, as well as recruited immune cells.

As HA inhibition with the repurposed compound hymecromone (4-MU) has been used successfully in human clinical trials (*32, 33*), and appears to be as an effective strategy in preclinical models of PSC, we have initiated a pilot clinical study in patients (NCT05295680). The findings reported here may inform that work and provide potential biomarkers to monitor treatment efficacy.

A limitation of our study is that the existing animal models may not faithfully replicate the pathogenesis of PSC. While both the DDC and Mdr2^-/-^ models are widely used better models are urgently needed that mirror the early changes in PSC in patients. Another issues is the lack of patient samples from earlier stages, as biopsies are very infrequently obtained. We have analyzed publicly available databases and found that CD44 and hyaluronan synthesis pathways are indeed enriched in PSC patients, compared to healthy controls. Furthermore in our study we did not perform detailed immune profiling, and although Kupffer cells do not appear to be an important source of HA (*15*), it is possible that T cell mediated local microenvironment changes can play a role.

In conclusion, we have identified an important role for CD44 as an mechanosensor for peribiliary ECM stiffness in early PSC caused by HMW-HA deposition, which leads to a reactive cholangiocyte phenotype with proliferation and proinflammatory cytokine production. Targeting HA synthesis and deposition early on promises to be a novel approach to mitigate fibroinflammatory responses in PSC.

## Materials and Methods

### Human samples

All human samples were de-identified and exempted (exemption 4) and were obtained from the Stanford Diabetes Research Center, Donor Network West (DNW), and the Stanford Tissue Bank. This was approved by Stanford University Institutional Review Board (IRB, 67378). Histology was evaluated by a hepatopathologist (GC) in a blinded manner.

### Mouse models

All animal experiments were performed in accordance with protocols approved by the Institutional Animal Care and Use Committee at Stanford University, and the Palo Alto Veteran’s Administration Hospital. Mice were maintained at 20–26 °C and 30–70% humidity and housed in standard cages under 12 hour light/12 hour dark cycles with ad libitum access to food and water unless otherwise specified.

A group of mice were then fed either a 0.1% DDC-containing diet (TD.07571, Teklad Diets) or regular chow (TD.00588, Teklad Diets) as a control for 48 or 96 hours. Mice were euthanized upon the completion of experiments and liver samples were collected for further experiments.

The OPN-iCreERT2 mice have a CD1-enriched background and were a kind gift from Lindsey Kennedy (U. of Indiana). CD44^fl/fl^ mice have a C57BL/6 background and were a kind gift from Dr. Paul Bollyky (Stanford University). The inducible CD44^BEC-KO^ mice were generated by crossing OPN-iCreERT2 with CD44^fl/fl^. The transgenic lines were backcrossed for several generations into C57BL/6. The transgenic allele was genotyped by PCR using the following primers: Osteopontin CreER genotyping primers: forward, GAAATTGCCCTTTTCCTTGC; reverse, AGGATCTGCACACAGACAGG. CD44 wild-type primers: forward, GCTGTTACTTGGTTTTCATGTGTGT; reverse, CTGAGGAATGAAAGCCAATGTTTTTCT. CD44 floxed primers: forward, TGCATGTGTGTGTAGGTTCTCATTT; reverse, GGGCCCGGTACCCAAT. In experiments, 8-week-old male CD44 ^BEC-KO^ and CD44^fl/fl^ littermate controls were used. Mice were injected intraperitoneally with tamoxifen (T5648, Sigma; stock concentration 20 mg/mL) at a dose of approximately 75 mg/kg body weight, or vehicle. Tamoxifen/vehicle were administered once every 24 hours for a total of 5 consecutive days, prior to starting the DDC diet.

FVB/N Mdr2^-/-^ mice were maintained in our breeding colony as was previously described (*34*). Four-week-old male FVB/N mice (wild-type or Mdr2^-/-^) received either regular drinking water or water supplemented with 4-MUG (ChemImpex) at 2 mg/mL concentration, corresponding to a daily dose of 10 mg per mouse, or vehicle. Mice were sacrificed at 8 weeks. Liver samples were collected for immunohistological and molecular analyses.

### Tissue immunofluorescence and immunohistochemistry

Paraffin-embedded tissue samples were cut into 5□µm sections, deparaffinized and rehydrated. For antigen retrieval, slides were boiled in citrate buffer (0.01□M, pH□6.0) using a microwave oven on high power for 5□min and cooled down to room temperature. After incubation in 3% aqueous H_2_O_2_ to quench the endogenous peroxidase, the sections were washed in PBST (PBS with 0.1% Tween-20, v/v) washing buffer, blocked with 3% BSA and 5% goat serum (EMD Millipore) diluted in PBST at room temperature for 1□h and incubated with primary antibodies diluted in the blocking buffer at 4□°C overnight. The slides were incubated in the appropriate secondary antibodies (Table 1).

**Table 1.**
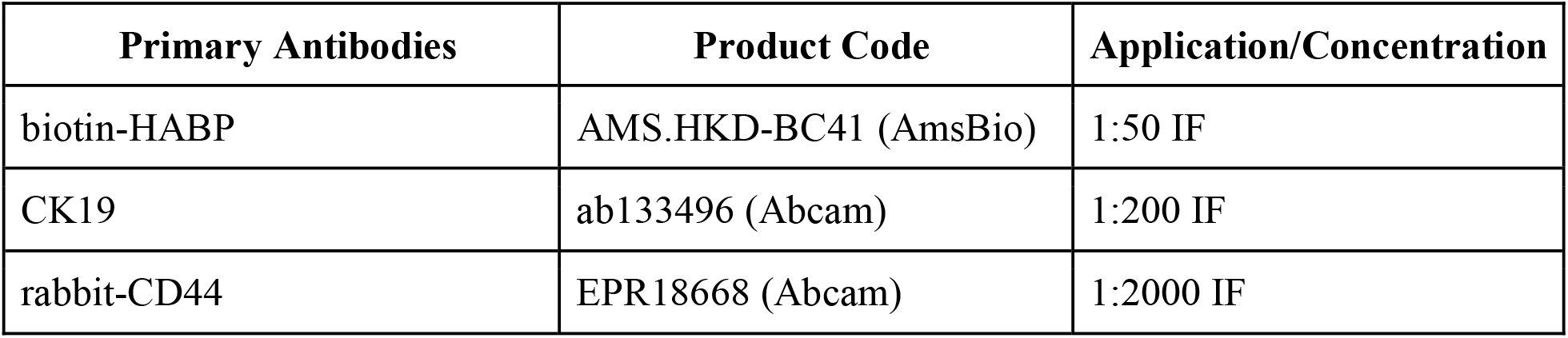
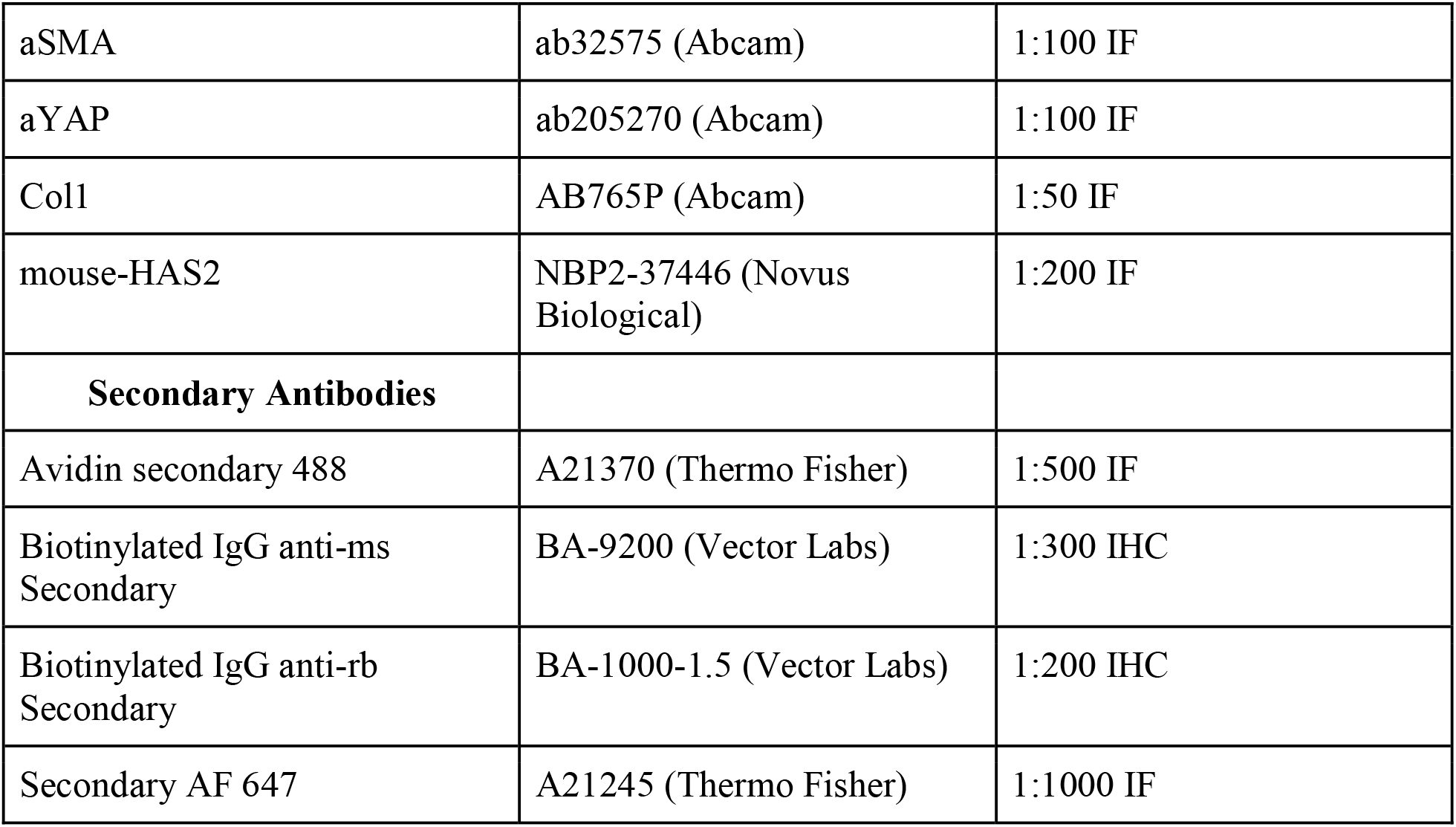
List of antibodies used for various experiments.

**Table 2.**
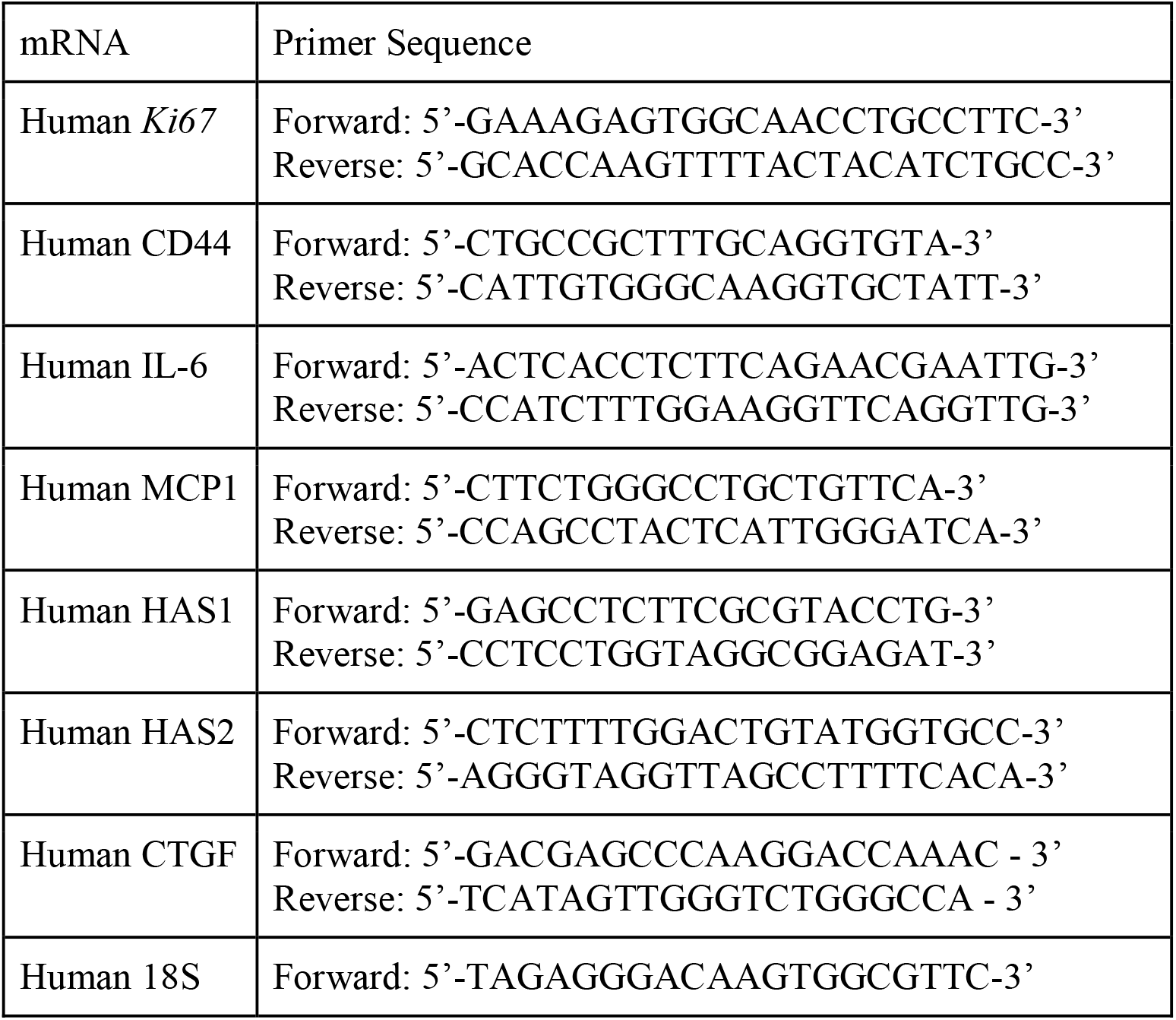
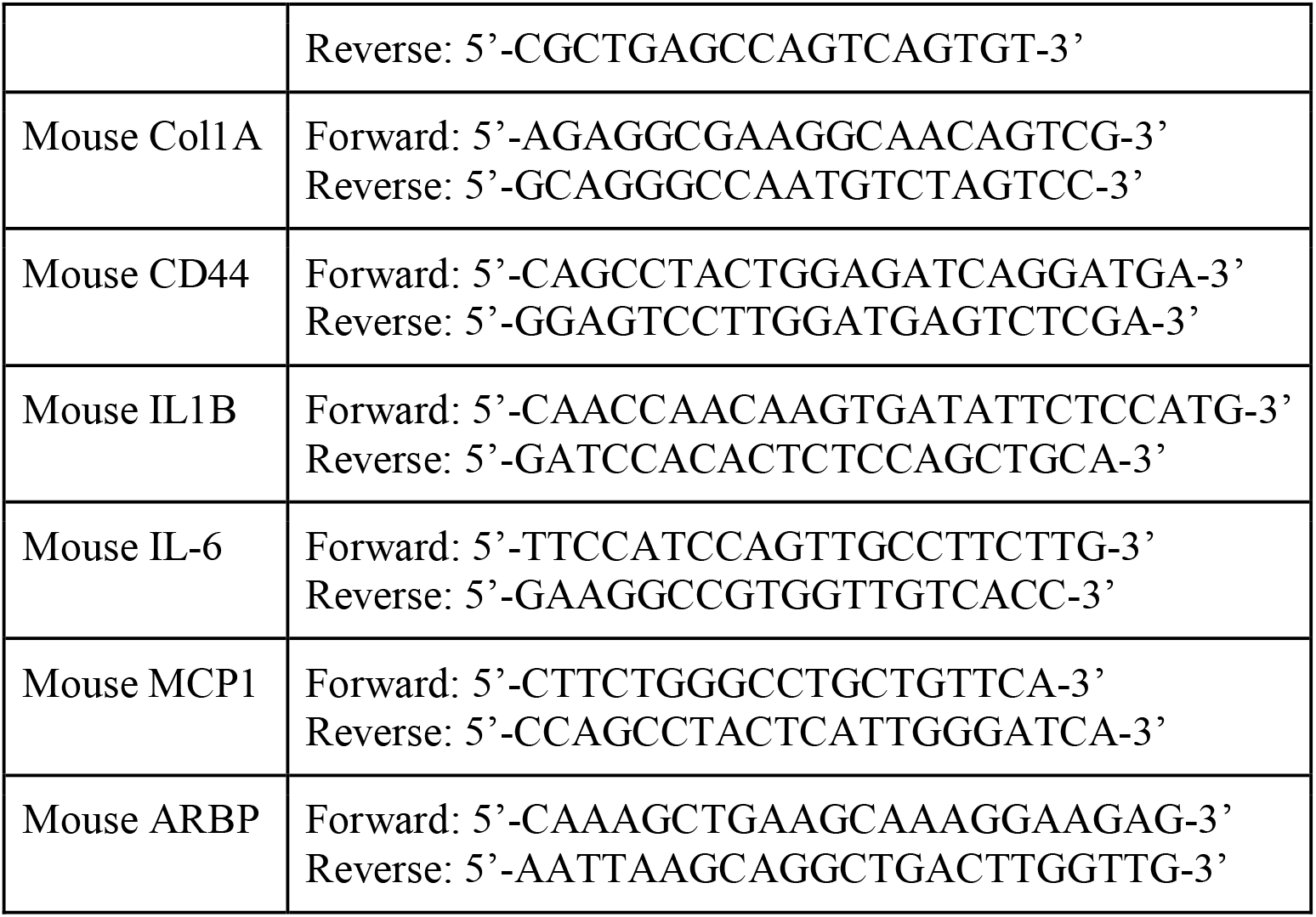
List of primer sequences for RT qPCR.

For immunohistochemistry slides were incubated with appropriate biotinylated secondary antibodies (Supplementary Table 1) for 1□h, and then processed according to the ABC Peroxidase Standard Staining Kit (Thermo Fisher Scientific) for 30□min. The slides were stained with 3,3′-diaminobenzidine (Abcam) and counterstained with hematoxylin (Thermo Fisher Scientific) for 45□s. The images were scanned using the Leica Aperio AT2 and Keyence BZ_X810 systems.

### Hydroxyproline assay

The liver tissue was denatured in 6 N HCl at 100 1C for 16 h, washed, and then resuspended in H_2_O. The tissue suspension was then mixed with chloramines-T (50 mM) and incubated at room temperature for 20 min. After the addition of perchloric acid (3.15 M, Sigma-Aldrich) and 20% p-dimethylaminobenzaldehyde (Sigma-Aldrich), the absorbance at 557 nm was measured. The hydroxyproline amount was calculated based on a standard curve generated from a series of hydroxyproline solutions with known concentrations. The data were expressed as microgram of hydroxyproline per gram of wet liver.

### Collagen imaging

Collagen fibers in mouse samples were analyzed by second harmonic generation (SHG) microscopy. All the samples were imaged using the Leica TCS SP5 multiphoton confocal microscope or the Leica Stellaris 8 DIVE upright confocal microscope. The excitation wavelength was tuned to 840□nm, and a 420□±□5□nm narrow band-pass emission controlled by a slit was used for detecting the SHG signal of collagen. The images were recorded using an inverted confocal laser-scanning microscope (Leica TCS SP8) equipped with a ×20 water-immersion objective for confocal reflection imaging. An Ar+ laser at 488□nm was used to illuminate the sample, and the reflected light was detected with photomultiplier tube (PMT) detectors. Scans were at 1,024□×□1,024 pixels, and all the images were taken 80–100 μm into the samples.

### Atomic force microscopy

Frozen liver tissue embedded in OCT compound were used (Sakura) that had been snap-frozen by direct immersion in liquid nitrogen and sectioned at 100 μm using a Leica CM1900-13 cryostat. Sections were pre-warmed in an oven to improve tissue adherence onto the glass slide, and OCT was dissolved in PBS for 5 min prior to measurements. Measurements were acquired on either a Bruker Resolve BioAFM or a JPK NanoWizard® V. Quantitative micromechanical mapping was performed using pre-calibrated spherical probes: Novascan-modified silicon nitride cantilevers (nominal spring constants k=0.01 and 0.05 N.m-1) bearing borosilicate spheres (nominal diameters 10 μm and 5μm, respectively), and Bruker SAA-SPH-5UM probes (5 μm radius, nominal k=0.21 N.m-1). Force maps (Force Volume mode) of 94.7 × 94.7 μm^2^ were collected around bile ducts in portal areas identified by Hoescht nuclear staining (1:1000 dilution, 20 min) under fluorescence microscopy (Zeiss). The sections were maintained at ambient temperature in PBS containing Halt™ Protease Inhibitor Cocktail (1:100) to limit proteolysis for the duration of the experiment. Force curves were recorded with an approach velocity of 1 μm.s-1 and a trigger force of 3 nN, with maximum indentation limited to <10% of local section thickness to minimize substrate effects and remain within the small-strain regime. Cantilever spring constants and optical level sensitivities were determined by the thermal tune method immediately before each session; nominal manufacturer values are reported above. Elastic (Young’s) moduli were obtained by baseline correction, contact-point identification, and fitting the loading segment of force-displacement curves to the Hertz model for a spherical indenter (F=4/3 E* R1/2 δ3/2), assuming a sample Poisson’s ratio of 0.5.

For the wild type DDC/chow cohort, reported values are the mean Young’s modulus measured from three bile duct/portal-region fields and three of central fields per mouse, averaged across n=5 or n=3 mice/group, respectively. For the cholangiocyte-specific CD44 KO cohort, 3 peribiliary areas/ mouse were sampled from n=4-5 mice; for each duct we averaged all force-displacement curves with that duct to produce a single duct-level Young’s modulus, and results were presented as median ± IQR of these duct means; as an alternative visualization all individual force-displacement curves were compiled into a single distribution and shown as a histogram. Data analyses were performed using MountainsLab® Premium V10 or JPK SPM Data Processing software and custom scripts in R (RStudio 2024.04.02).

### Collagen 3D hydrogel formation

The alginate (Provona UP VLVG, NovaMatrix) was transferred to a 1.5□ml Eppendorf tube (a polymer tube) and kept on ice. Telopeptide intact collagen I (rat tail, Corning) was neutralized (pH ∼7.4) using 10X DMEM (Gibco) and 1 M NaOH (Merck) on ice. Calcium sulfate was added to 1□ml Luer lock syringes (Cole-Parmer) and stored on ice to maintain the constant Young’s moduli of the substrates. The polymer mixes were divided into individual 1 ml Luer lock syringes (polymer syringes) and placed onto ice as well. The polymer syringe was linked to the calcium sulfate syringe to create gels. The two solutions were quickly combined using 30 pumps on the syringe handles, and the resulting mixture was placed into a well of an eight-well Lab-Tek plate (Thermo Fisher Scientific) and was allowed to gel before adding media and placing it in the incubator.

The collagen–HA gels were prepared by neutralizing rat tail collagen I (Corning) with 1 N NaOH and mixing to a final concentration of 2.5 mg/mL with low- and high-molecular-weight HA at a concentration of 1 mg/mL (100 kDa and 1.5 MDa sodium hyaluronate (R&D systems), respectively). The solution was directly deposited onto the lower plate of the rheometer for rheological testing.

### Hydrogel shear rheology

Rheology experiments were performed using a stress controlled AR2000EX rheometer (TA Instruments). IPNs were directly deposited onto the lower 25 mm Peltier plate for rheology testing. To prevent dehydration of the hydrogel, mineral oil (Sigma-Aldrich) was applied to the gel’s edges. The storage and loss moduli had equilibrated by the time the time sweep was done, which was at 1□rad□s−1, 37□°C and 1% strain.

### 3D cell culture

H-69 cells (cholangiocyte cell line) were gifted from Dr. Greg Gores (Mayo Clinic). DMEM/F12 was the base medium and it was supplemented with 5 ug/mL insulin, 24.3 ug/mL Adenine, 1 ug/mL Epinephrine, 0.0013 ug/mL triiodothyronine, 10.5 EGF, 0.4 ug/mL Hydrocortisone, and 5 ug/mL Transferrin. Cells were passaged 2-3 times a week. For the 3D hydrogel experiments, H69 cells were incorporated into the hydrogels with tunable stiffness. Briefly, cells dissociated using 0.05% trypsin/EDTA, washed, centrifuged, and resuspended in serum-free medium. Cell density was determined using a Vi-Cell Coulter counter (Beckman Coulter). Type I collagen was combined with low molecular weight alginate, after which cells were incorporated into the polymer mixture and loaded into a pre-cooled syringe. The mixture was vigorously combined with a CaSO□solution and dispensed into chambered cover glass wells (Lab-Tek). The final concentrations of collagen and alginate were 1.6 mg/ml and 4.8 mg/ml, respectively. CaSO□concentrations of 10 mM, and 35 mM were used to generate IPNs with defined stiffness values of 1 kPa and 2.5 kPa, respectively. The final cell density was 3 × 10□cells per ml. The hydrogels were incubated for 60 minutes at 37 °C in a humidified atmosphere containing 5% CO□, 10% FBS containing DMEM was added. Culture medium was replaced after 24 hours. After 48 hours, cells were harvested for further processing. To inhibit integrin β1, monoclonal integrin-β1-blocking antibody (Abcam, P5D2, 1.5□μg/ml) and/or CD44 blocking antibody (1.5 μg/mL, Thermo Fisher) were diluted in 250 μL of media and added to the cells for 1 hour before encapsulating the cells in the hydrogels. IgG nonspecific antibody (Sigma-Aldrich, I5381) was used as a negative control.

### Proximity ligation assay (PLA)

The Duolink proximity ligation assay kit (Sigma-Aldrich) was used to determine the interaction between CD44 and integrin β1 in H69 cells. The anti-CD44 and anti-integrin β1 were used as the primary antibodies, bound to a pair of oligonucleotide-labelled secondary antibodies (PLA probes), the hybridizing connector oligos joined the PLA probes if they were close and the ligase created the DNA template needed for rolling-circle amplification. Labelled oligos hybridized to the complementary sequences in the amplicon and produced discrete red fluorescent signals that could be seen by confocal microscopy (Leica TCS SP8l). NIH Image J (v.1.53t) software was used to count the signal, and the average counts were used for the plot.

### Transfection experiments

H69 cells were maintained in complete growth medium, consisting of a 1:1 (v/v) mixture of DMEM and Ham’s F-12 supplemented with a growth factor cocktail including insulin, adenine, epinephrine, triiodothyronine-transferrin, epidermal growth factor, and hydrocortisone. Cells were cultured at 37°C in a humidified 5% CO2 atmosphere and seeded into 6-well or 24-well plates until reaching approximately 80% confluence. To investigate CD44 signaling, H69 cells were transiently transfected with 3 μg of plasmid DNA encoding mutant variants (CD44R41A, or CD44v2-v10, as control). Transfections were performed using Lipofectamine 3000 (Invitrogen) in Opti-MEM Reduced Serum Medium for 48 hours. Transfection efficiency was monitored by parallel delivery of an EGFP-N1 reporter plasmid and quantified via fluorescence microscopy scoring of EGFP-positive cells. Following transfection and culture under differential mechanical conditions (soft vs. stiff substrates) for 24-48 hours, total RNA was extracted for gene expression analysis.

### RT-qPCR

Total mRNA was extracted from snap-frozen liver tissues and cells in hydrogels using the Trizol method (Invitrogen). Complementary cDNA was created from an identical amount of RNA using the iScript cDNA synthesis kit (Bio-Rad). The PowerUp SYBR Green PCR Master Mix (Applied Biosystems) was used for quantitative PCR with reverse transcription (RT–qPCR) on the 7900HT machine (Applied Biosystems), and the data were evaluated using the 2−*C*t technique. As an endogenous control, *Arbp* (also known as *Rplp0*) or *18S* were used to standardize the data. A list of the primer sequences used in this study is provided in Supplementary Table 2.

### Bioinformatic analyses of single-cell RNA sequencing (scRNA-Seq) in healthy and PSC patient livers

#### Data import and preprocessing

Single-cell RNA-seq data (GSE247128) from normal and PSC livers were downloaded as an AnnData .h5ad object. The .h5ad file was read in Python (Scanpy) and exported to 10x Genomics–style sparse matrices using write_10x_mtx, yielding matrix.mtx, features.tsv, barcodes.tsv, and a cell-level metadata table (cell_metadata.csv). These files were imported into R (R Studio 2024.04.2) and Seurat (v4.4) by constructing a count matrix with ReadMtx and creating a Seurat object with CreateSeuratObject, after which the original metadata (donor_id, disease, and cell_type annotations) were reattached using AddMetaData. Cells from primary biliary cholangitis donors were excluded, and the analysis was restricted to cells annotated as “normal” or “primary sclerosing cholangitis” in the original metadata.

#### Quality control and filtering

For each cell we quantified the number of detected genes (nFeature_RNA), total UMI counts (nCount_RNA), and the fraction of UMIs mapping to mitochondrial genes (percent.mt, defined using PercentageFeatureSet with prefix “MT-”). Cells were retained if they satisfied all of the following criteria: nFeature_RNA > 250, nFeature_RNA < 7,500, nCount_RNA > 500, and percent.mt < 20. These thresholds removed putative empty droplets, low-complexity cells, and cells with high mitochondrial content while retaining most high-quality profiles from both normal and PSC donors. Following QC filtering we retained a total of 24,210 normal cells from 6 healthy donors and 46,764 cells from 8 PSC donors that were subjected to downstream analysis.

#### Normalization, feature selection, and dimensionality reduction

All analyses were performed on the RNA assay. Raw counts were log-normalized using Seurat’s NormalizeData with default parameters, and highly variable genes were identified with FindVariableFeatures (3,000 features). Gene expression was scaled with ScaleData, regressing out library size (nCount_RNA) and mitochondrial percentage (percent.mt) to mitigate technical effects. Principal component analysis was performed with RunPCA (50 PCs), and the number of informative components was selected based on the elbow plot (first 30 PCs used in downstream steps).

#### Batch correction, clustering, and cell type annotation

To reduce donor-specific batch effects, we applied Harmony integration on the PCA embedding with RunHarmony, using donor_id as the grouping variable and the first 30 PCs as input dimensions. The Harmony embedding was used to construct a shared nearest neighbor graph (FindNeighbors) and to compute a UMAP representation (RunUMAP). Clusters were identified using Seurat’s Louvain clustering (FindClusters, resolution 0.4). Cell types were assigned by mapping the original cell type labels from the metadata onto the integrated object and inspecting canonical marker expression on the UMAP, yielding major liver populations including intrahepatic cholangiocytes.

#### Intrahepatic cholangiocyte subsets and CD44 PSC cells

For focused analyses, we subset the integrated object to intrahepatic cholangiocytes re-normalized this subset, re-computed variable features, and re-scaled the data (regressing out nCount_RNA and percent.mt), followed by PCA (20 PCs) and UMAP computed directly on the PCA space (15 dimensions). The resulting embedding contained both normal and PSC cholangiocytes, disease labels were recoded into a human-readable factor (disease_pretty) and used to visualize normal versus PSC cholangiocytes on a shared UMAP. CD44 expression within cholangiocytes was examined by UMAP FeaturePlots of CD44 alongside cholangiocyte markers (KRT19, KRT7, EPCAM). CD44-positive cholangiocytes were defined as cells with detectable CD44 transcript (CD44 > 0), and a CD44 status metadata factor (“CD44□” vs “CD44□”) was created accordingly. The proportion of CD44 cholangiocytes per disease state was calculated with dplyr and visualized as a 100% stacked bar plot.

#### Differential expression and pathway analysis in PSC cholangiocytes

To compare transcriptional programs between CD44□ and CD44□ cholangiocytes in PSC, we restricted the cholangiocyte subset to PSC donors. Within this PSC cholangiocyte subset, we set the cell identities to CD44 status and performed differential expression using Seurat’s FindMarkers (Wilcoxon rank-sum test, ident.1 = “CD44+”, ident.2 = “CD44-”, logfc.threshold = 0, min.pct = 0.05). For gene-set enrichment, differentially expressed genes were ranked by a composite metric combining effect size and significance (average log2 fold change × –log10 p-value). This ranked list was used as input to fgsea with Hallmark (category H) gene sets obtained from msigdbr (Homo sapiens). Enrichment results were filtered by adjusted p-value (FDR < 0.25), and the top up- and down-regulated Hallmark pathways were visualized as a dot plot annotated with normalized enrichment score (NES) and –log10(FDR).

#### Mechanotransduction and hyaluronan biosynthesis module scores

To interrogate mechanotransduction programs, we defined a custom integrin/focal adhesion mechanotransduction gene set enriched for CD44/ITGB1-linked signaling. Within PSC cholangiocytes, we extracted the log-normalized expression matrix (GetAssayData(assay = “RNA”, layer = “data”)) and ranked gene expression per cell using AUCell. The mechanotransduction gene set (restricted to genes present in the matrix) was passed to AUCell_calcAUC to obtain per-cell AUC scores. The resulting “Integrin_FA_mech_AUC” scores were stored in the chol_psc metadata and visualized as (i) violin/boxplots comparing CD44□ vs CD44□ PSC cholangiocytes and (ii) UMAP FeaturePlots overlaid on the cholangiocyte embedding. Differences between CD44□ and CD44□ groups were assessed using a Wilcoxon rank-sum test on the AUCell scores. To quantify hyaluronan synthesis using a non-custom gene set, we retrieved Gene Ontology Biological Process (GO:BP) gene sets from msigdbr (C5:GO:BP, Homo sapiens) and extracted the GO term “hyaluronan biosynthetic process” (GO:0030213; HYALURONAN_BIOSYNTHETIC_PROCESS). The corresponding gene symbols, intersected with the expressed genes in PSC cholangiocytes, defined a “GO_HA_BIOSYNTHESIS” gene set. Per-cell AUCell scores for this GO term were computed in the same manner as for the mechanotransduction module and stored as “GO_HA_Biosynthesis_AUC”. AUCell scores were compared between CD44□ and CD44□ PSC cholangiocytes by Wilcoxon test and visualized by violin/boxplots and overlaid on the cholangiocyte UMAP to show their spatial relationship to CD44 expression.

#### Statistical analyses

At least three biological replicates were performed for all in vivo and in vitro experiments. Data are presented as mean□±□s.e.m. Statistical analyses were performed using GraphPad Prism v.11.0.0 (GraphPad Software). Two-tailed unpaired t-tests and one-way ANOVA with Tukey tests were used to analyze data with a normal distribution. Data with non-equal variance standard distributions were analyzed using Welch’s t test. P□<□0.05 was considered to be statistically significant.

## Acknowledgments

Patient samples were obtained using the services of the Diabetes Clinical and Translational Core facility of the Stanford Diabetes Research Center which is supported by the National Institute of Diabetes and Digestive and Kidney Diseases of the National Institutes of Health under Award Number P30DK116074. Research organization/authors hereby express their thanks for the cooperation of Donor Network West and all of the organ and tissue donors and their families, for giving the gift of life and the gift of knowledge, by their generous donation.

Part of this work was performed at the Stanford Nano Shared Facilities (SNSF), supported by the National Science Foundation under award ECCS-2026822. Part of this work was performed at Stanford Cell Sciences Imaging Facility (CSIF, RRID:SCR_017787), supported by the Award Number 1S10OD010580-01A1, S10RR02557401, and 1S10OD032300-01 from the National Center for Research Resources (NCRR). Its contents are solely the responsibility of the authors and do not necessarily represent the official views of the NCRR or the National Institutes of Health. We are grateful to Dr. Nadia Makarova (Stanford SNSF) for the AFM, and to Gordon X. Wang (Wu Tsai Neuroscience Institute’s Neuroscience Microscopy Service) for assistance with imaging.

This research was supported by funding by the VA I01 BX002418, the NIH grant R01CA277710, and Stanford IMA Award to NJT. The study was also funded in part by the Boehringer Ingelheim Fund to AF.

---

**Supplementary Figure 1.**
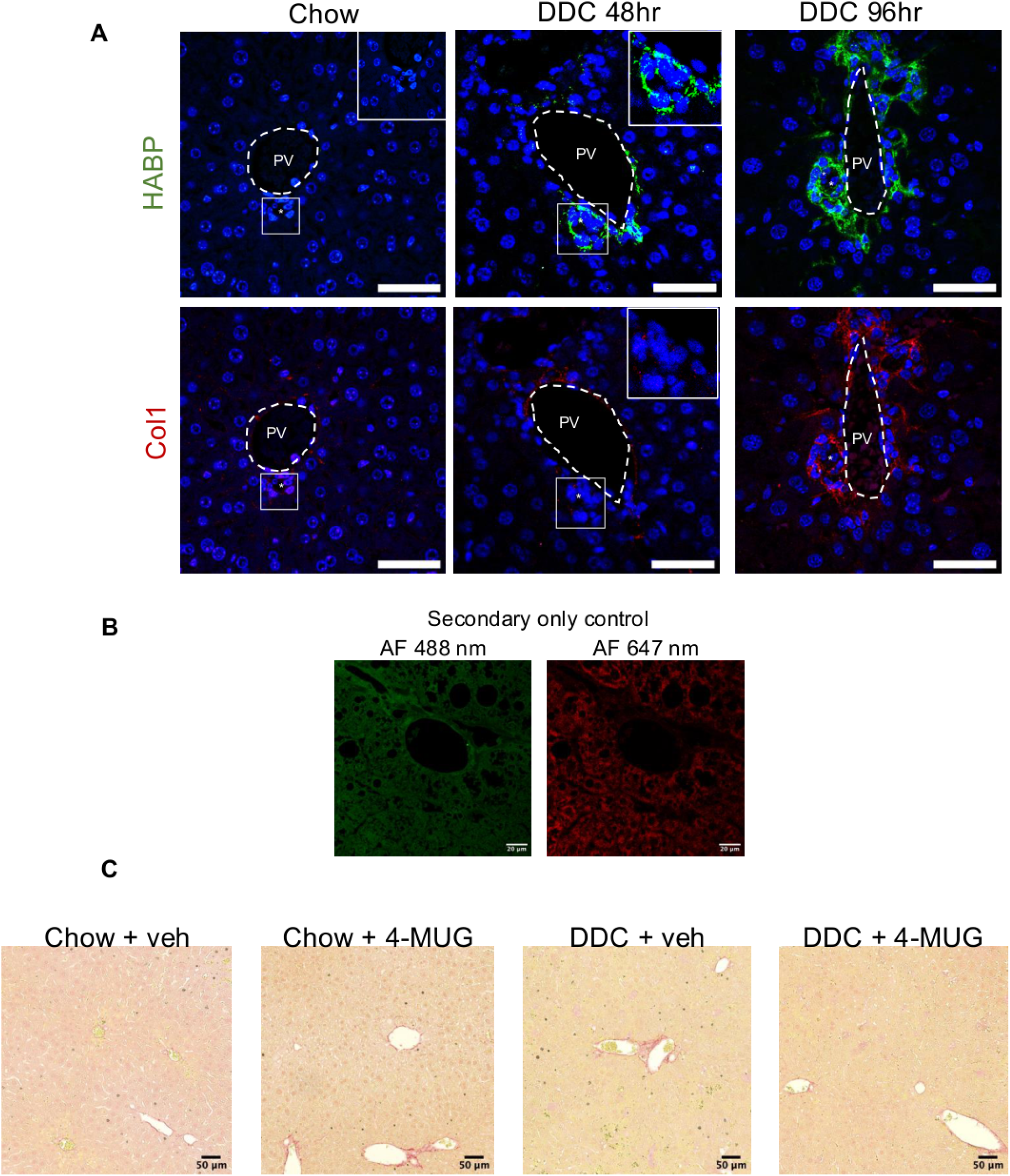
HABP, collagen 1 signals and picrosirius red signals across dietary and treatment conditions. **(A)** Representative confocal immunofluorescence (IF) images of HABP and collagen 1 individual channel signals, after 48 and 96 h on DDC diet, corresponding to the merged images in Fig. 1B, middle panel. **(B)** Secondary-only antibody controls for AF 488 nm (green) and AF 647 nm (red) channels used in IF staining. Scale bars = 20 µm. **(C)** Representative picrosirius red staining from vehicle or 4-MUG-treated mice. Scale bars = 50 µm.

**Supplementary Figure 2.**
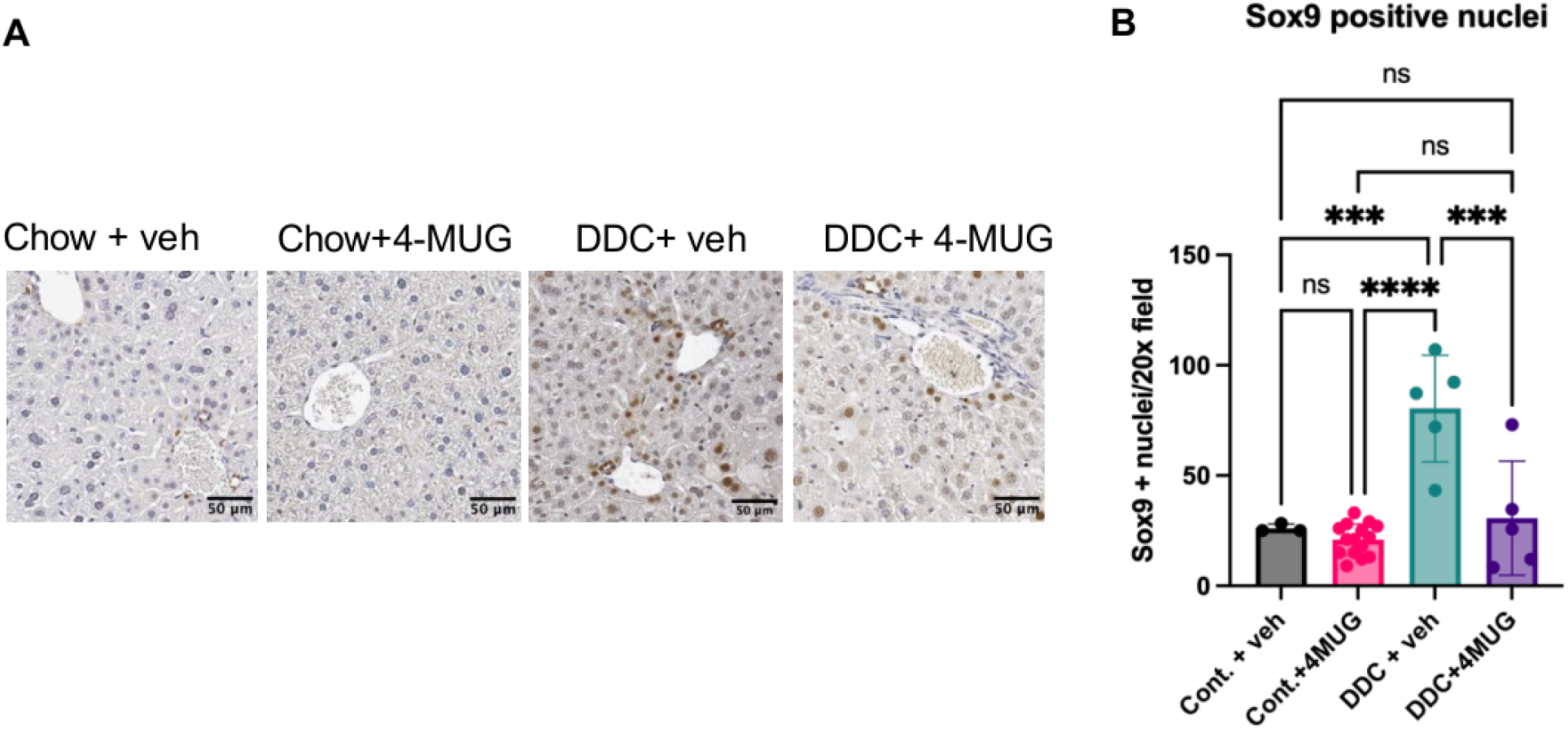
4-MUG treatment reduces DDC-induced Sox9 + cell expansion in the liver. **(A)** Representative immunohistochemistry for Sox9 (brown, nuclear) in liver sections from vehicle or 4-MUG-treated mice. Scale bars = 50 µm. **(B)** Quantification of Sox9-positive nuclei per 20× field across conditions. ***P < 0.005, ****P < 0.001, n=3-15, mean±SEM, one-way ANOVA.

**Supplementary Figure 3.**
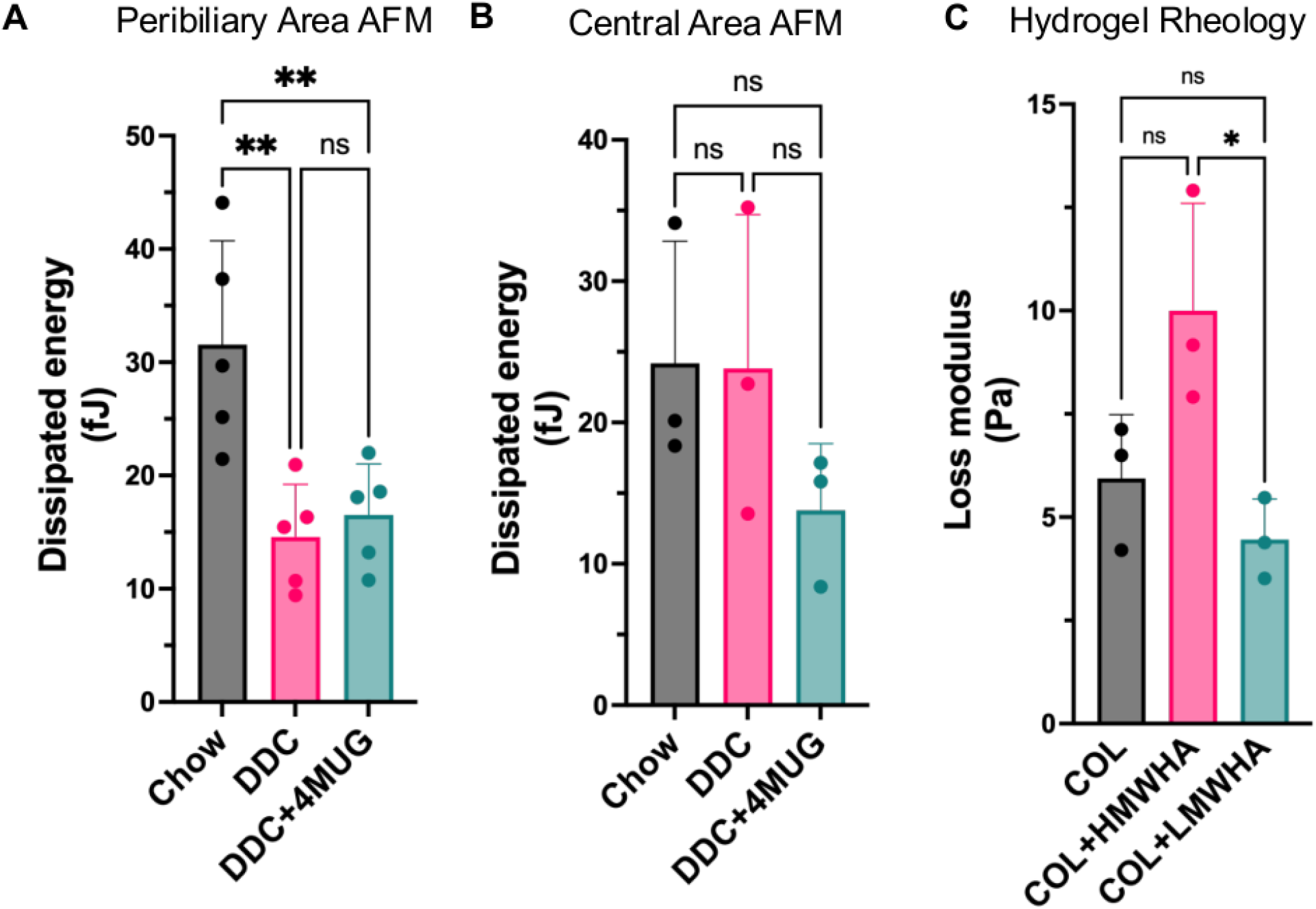
Mechanical characterization of liver tissues by AFM and hydrogel rheology. **(A)** Atomic force microscopy (AFM) measurements of dissipated energy (fJ) in the peribiliary areas of liver sections from mice on chow, or DDC diet, treated with vehicle or 4-MUG. **p < 0.01, ns: non-significant, n=5, mean±SEM, one-way ANOVA. **(b)** AFM dissipated energy measurements in the central, zone 3 regions. n=3, mean±SEM, one-way ANOVA. **(c)** Oscillatory rheology of collagen-based hydrogels measuring loss modulus (Pa) for collagen alone (COL), collagen supplemented with high-molecular-weight hyaluronic acid (COL + HMWHA), and collagen supplemented with low-molecular-weight hyaluronic acid (COL + LMWHA). Data are presented as mean ± SD with individual data points shown. ns = not significant, n=3, mean±SEM, one-way ANOVA.

**Supplementary Figure 4.**
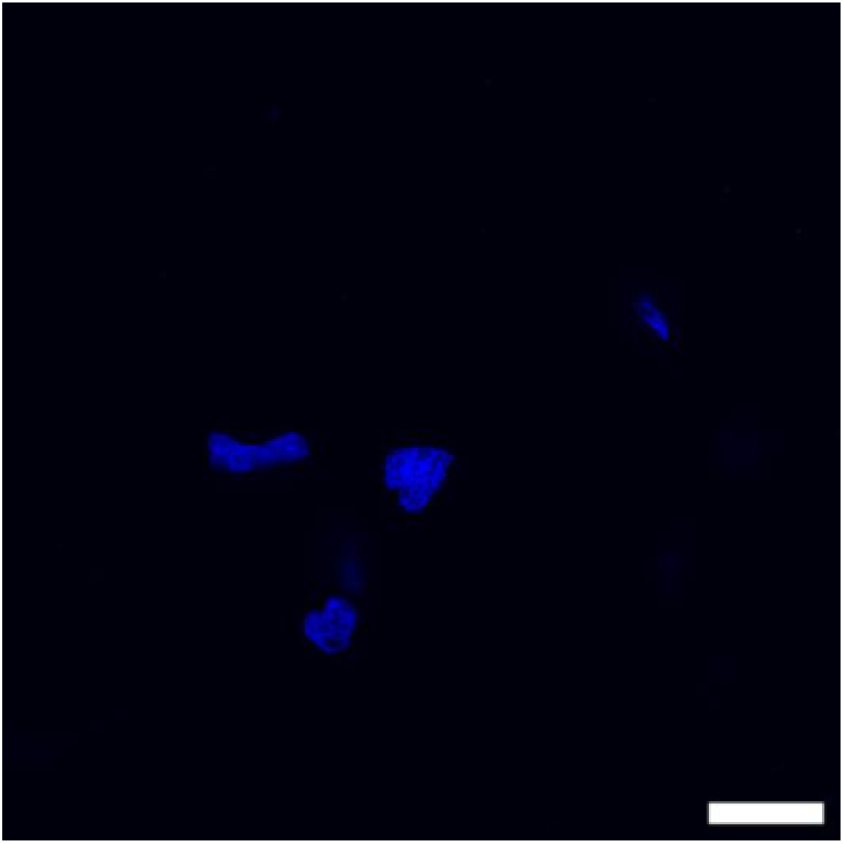
**Suppl. Fig. 4**. No antibody control for the proximity ligation assays Scale bar=10μm.

**Supplementary Figure 5.**
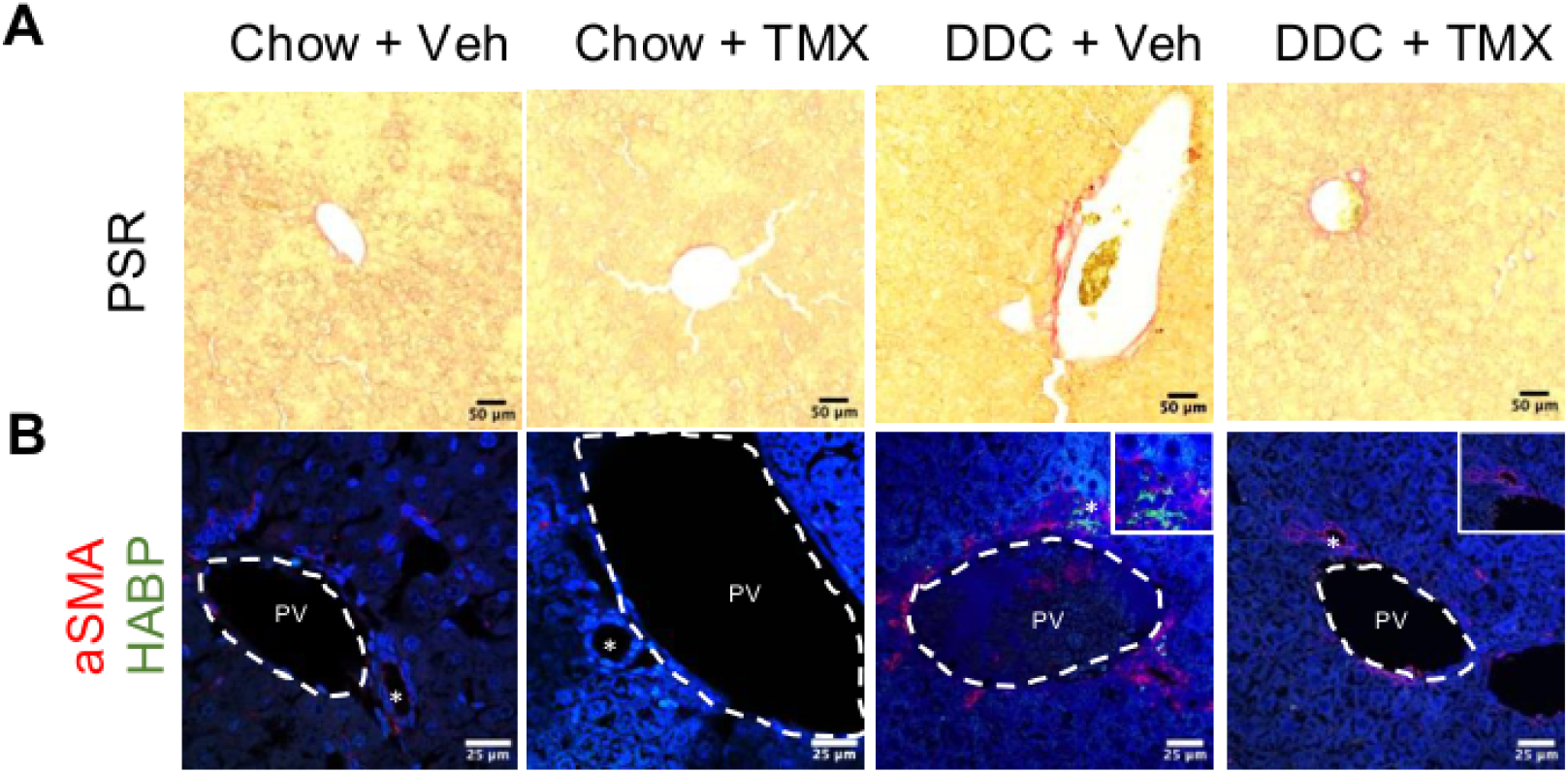
Picrosirius red and aSMA immunofluorescence at 48 hours on chow or DDC diet, in vehicle or tamoxifen-injected mice. **(A)** Representative images of Picrosirius Red (PSR) staining the different groups. Scale bars = 50 µm. **(B)** Immunofluorescence images, and confocal microscopy for α-smooth muscle actin (αSMA, red), and hyaluronic acid binding protein (HABP, green), with nuclei counterstained in blue (DAPI). Boxed areas are enlarged images of the bile ducts. PV: portal vein. Tmx: Tamoxifen; Scale bars = 25 µm.

**Supplementary Figure 6.**
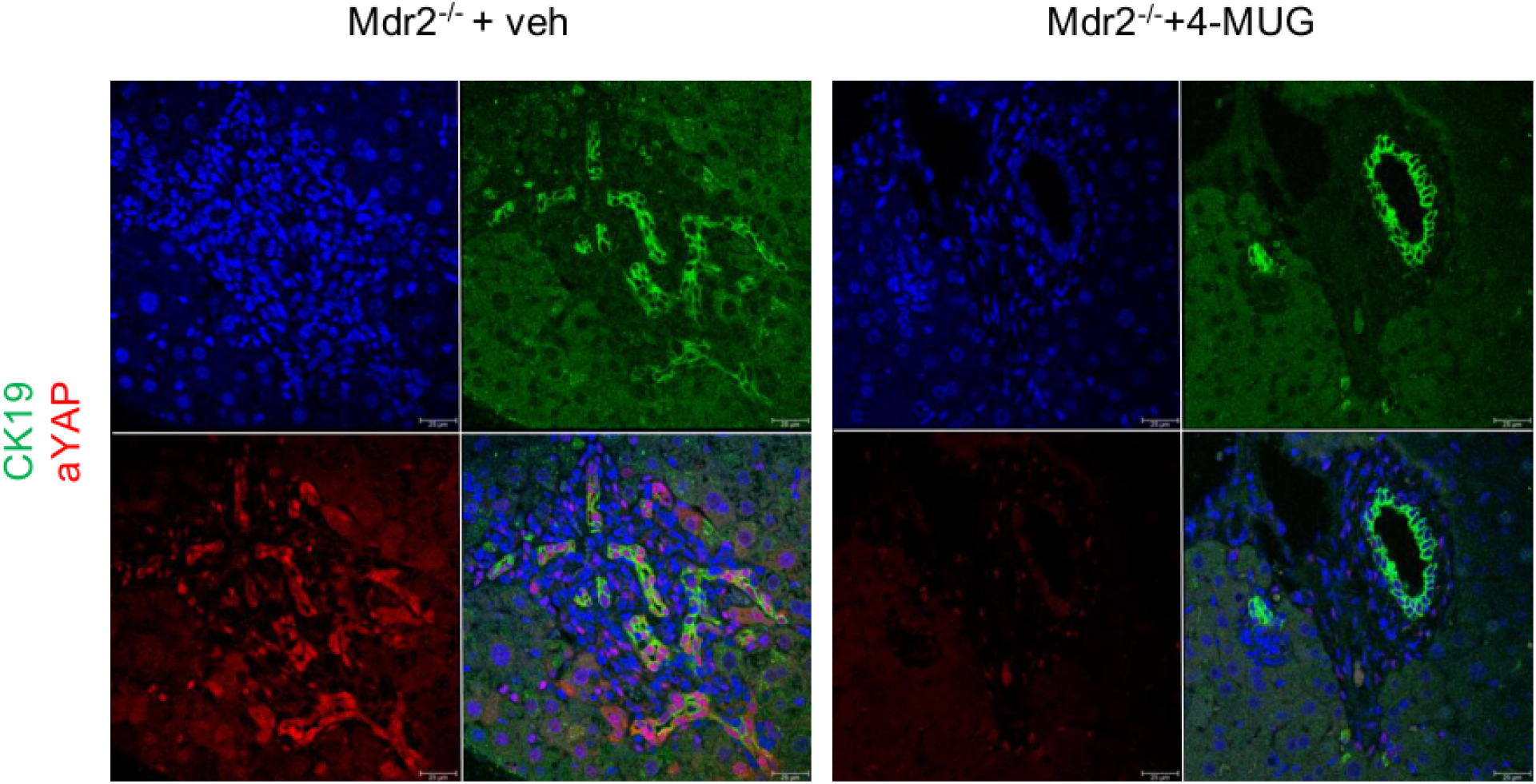
4-MUG treatment reduces YAP activation in CK19-positive ductal cells in Mdr2. □**/**□**mice.** Representative confocal immunofluorescence images of liver sections from vehicle or 4-MUG treated mice co-stained for CK19 (green), and active YAP antibody (aYAP, red), with nuclei counterstained in blue (DAPI). Each condition is shown as an individual channel (top row: DAPI and CK19; bottom row: aYAP and merged overlay). Scale bars = 25 µm.

**Supplementary Figure 7.**
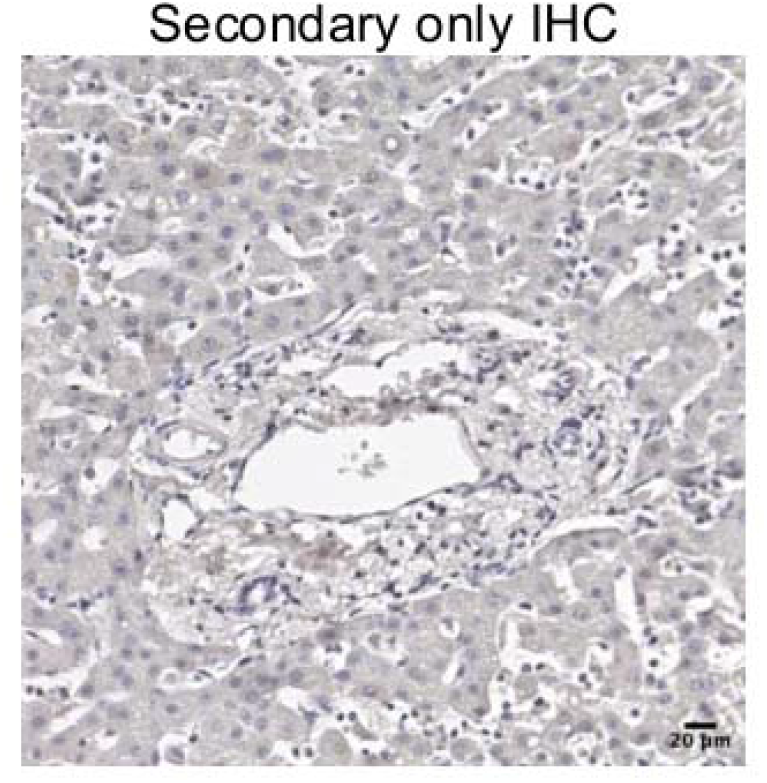
Secondary antibody-only control for immunohistochemistry. Representative image of a liver section processed with secondary antibody alone. Scale bar = 20 µm.

